# Chromatin profiling identifies transcriptional readthrough as a conserved mechanism for piRNA cluster biogenesis in mosquitoes

**DOI:** 10.1101/2022.08.22.504762

**Authors:** Jieqiong Qu, Valerie Betting, Ruben van Iterson, Florence M. Kwaschik, Ronald P. van Rij

## Abstract

The piRNA pathway in mosquitoes differs substantially from other model organisms, with an expanded PIWI gene family and functions in antiviral defense. Here, we defined core piRNA clusters as small RNA source loci that showed ubiquitous expression in both somatic and germline tissues. These core piRNA clusters were enriched for non-retroviral endogenous viral elements (nrEVEs) in antisense orientation and depended on key biogenesis factors, Nxf1, Veneno, Tejas, Yb, and Shutdown. Combined transcriptome and chromatin state analyses identified transcriptional readthrough as a conserved mechanism for piRNA cluster biogenesis in *Aedes aegypti*, *Aedes albopictus*, *Culex quinquefasciatus*, and *Anopheles gambiae*. Comparative analyses between two *Aedes* mosquitoes suggested that piRNA clusters function as traps for nrEVEs, allowing adaptation to environmental challenges such as virus infection. Our systematic transcriptome and chromatin state analyses lay the foundation for studies of gene regulation, genome evolution and piRNA functions in these important vector species.

**Graphical abstract:** 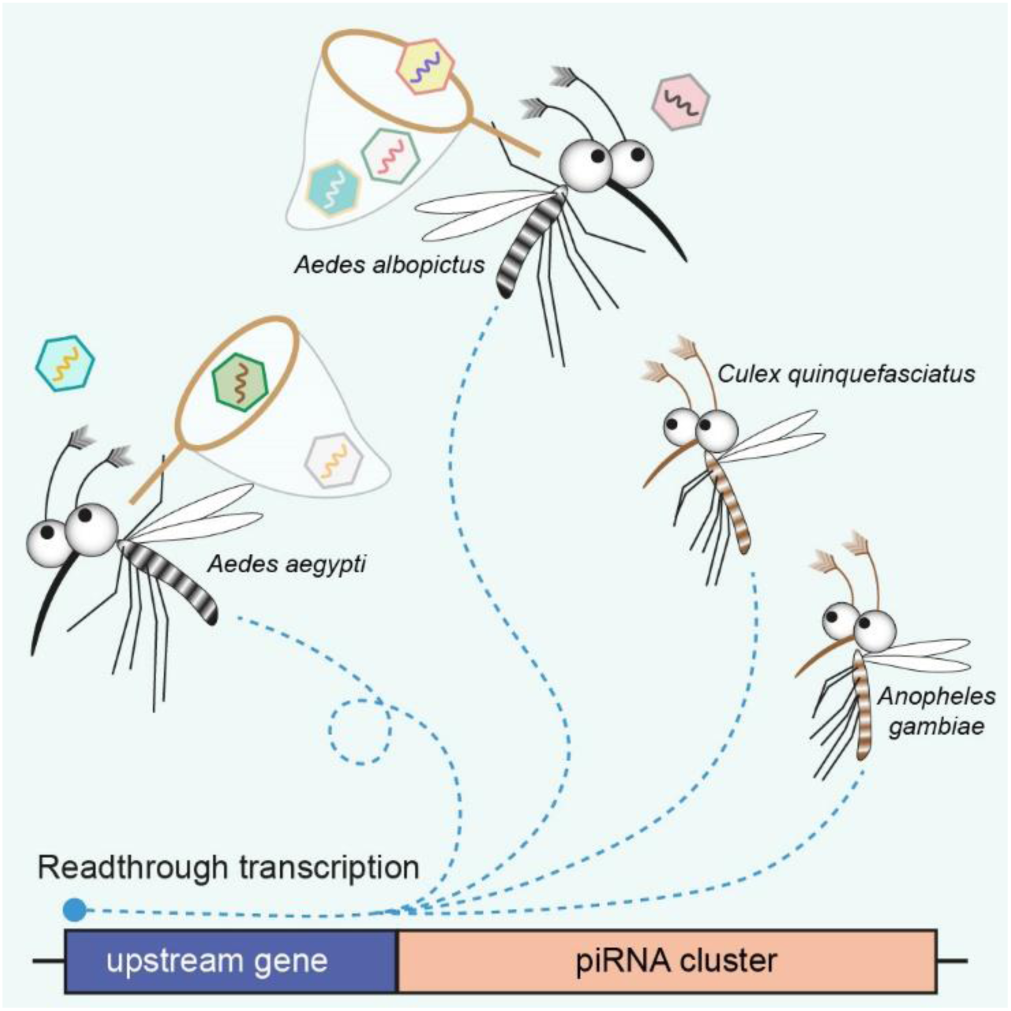

**Highlights:**

- Core piRNA clusters showed ubiquitous expression in both somatic and germline tissues in four vector mosquitoes.
- Chromatin profiling identifies transcriptional readthrough as a conserved mechanism for piRNA biogenesis.
- Biogenesis of cluster-derived piRNAs depends on key factors, Nxf1, Veneno, Tejas, Yb, and Shutdown.
- piRNA clusters function as traps for viral elements downstream of conserved set of genes in *Aedes* mosquitoes

## INTRODUCTION

*Aedes* mosquitoes transmit arthropod-borne (arbo) viruses including dengue, Zika, and chikungunya virus (Souza-Neto et al., 2019), posing significant health and economic impact (Chibueze et al., 2017; Diagne et al., 2021; Weaver and Reisen, 2010). A female mosquito acquires arboviruses through an infectious blood meal, after which viruses establish a systemic infection. Albeit permissive to systemic infection, mosquitoes have evolved efficient antiviral strategies to limit viral replication to non-pathogenic levels (Cheng et al., 2016; Lee et al., 2019), such as the small interfering (si)RNA pathway (Blair, 2011; Bronkhorst and van Rij, 2014). More recent studies have revealed that another class of small RNAs, PIWI-interacting RNAs (piRNAs), play a role in antiviral defense as well (Morazzani et al., 2012; Schnettler et al., 2013; Suzuki et al., 2020; Tassetto et al., 2019; Varjak et al., 2017).

The piRNA pathway is evolutionarily conserved in animals but has been most extensively studied in the ovary of *Drosophila melanogaster*. In fruit flies, the majority of piRNAs originate from discrete genomic loci, termed piRNA clusters, that are especially rich in transposable elements (TEs) (Brennecke et al., 2007). The resulting piRNAs silence TEs at the transcriptional and post-transcriptional levels to maintain genome integrity (Li et al., 2009a; Saito et al., 2006; Vagin et al., 2006). Two types of piRNA clusters exist in *D. melanogaster*, uni-strand and dual-strand clusters, which produce piRNAs from one or both genomic strands, respectively (Brennecke *et al*., 2007). In somatic follicle cells, piRNAs are exclusively produced from uni-strand piRNA clusters, such as the *flamenco* (*flam*) locus (Brennecke *et al*., 2007; Goriaux et al., 2014; Malone et al., 2009). Similar to protein-coding genes, *flam* is transcribed by RNA polymerase II and the resulting transcripts are regularly spliced, capped, and polyadenylated. The piRNA precursor transcript is subsequently exported by Nxf1/Nxt1 to Yb-bodies where primary piRNA biogenesis occurs (Dennis et al., 2016; Goriaux *et al*., 2014; Li *et al*., 2009a; Qi et al., 2011). In germline cells, piRNAs are predominantly produced from dual-strand piRNA clusters that are marked by the repressive histone mark H3K9me3. Dual-strand clusters require a heterochromatin-dependent transcription machinery involving the Rhino-Deadlock-Cutoff (RDC) complex and Moonshiner for transcription while inhibiting splicing, capping, and polyadenylation (Andersen et al., 2017; Chang et al., 2019; Chen et al., 2016; Klattenhoff et al., 2009; Le Thomas et al., 2014; Mohn et al., 2014; Pane et al., 2011; Zhang et al., 2014). A specialized nuclear export complex, consisting of an ovary-specific heterodimeric mRNA export receptor Nxf3-Nxt1, Bootlegger, and the TREX component UAP56, is recruited for export of the piRNA cluster transcript to the nuage where piRNA biogenesis factors accumulate (ElMaghraby et al., 2019; Kneuss et al., 2019; Mohn *et al*., 2014; Pek et al., 2012; Zhang et al., 2012). In the nuage, piRNA biogenesis occurs in a feed-forward loop, called ping-pong amplification, involving two PIWI proteins, Aubergine and Ago3 (Brennecke *et al*., 2007; Lim and Kai, 2007; Saito *et al*., 2006; Vagin *et al*., 2006).

While hallmark features are conserved (Betting et al., 2021; Halbach et al., 2020; Joosten et al., 2019; Joosten et al., 2021a; Miesen et al., 2015), the piRNA pathway in *Aedes* mosquitoes differs significantly from *D. melanogaster* in terms of an expanded PIWI gene family and ubiquitously expressed piRNAs in somatic tissues (Akbari et al., 2013; Campbell et al., 2008; Ma et al., 2021; Marconcini et al., 2019; Miesen et al., 2016b). The majority of piRNAs in *D. melanogaster* are generated from TE-rich regions despite an overall low genomic TE content (Bergman et al., 2006; Brennecke *et al*., 2007; Mérel et al., 2020), whereas it is the opposite for *Aedes* mosquitoes (Arensburger et al., 2011). Moreover, the mosquito piRNA pathway processes a broader repertoire of substrates including transcripts from protein-coding genes and satellite repeats as well as cytoplasmic viral RNA, and, thus, has important functions beyond transposon silencing (Betting *et al*., 2021; Girardi et al., 2017; Halbach *et al*., 2020; Joosten *et al*., 2019; Miesen *et al*., 2015; Miesen et al., 2016a; Morazzani *et al*., 2012; Vodovar et al., 2012).

The genomes of main arbovirus vectors, *Aedes aegypti* and *Aedes albopictus*, contain sequences of non-retroviral RNA viruses (non-retroviral endogenous elements, nrEVEs) from several families including the *Flaviviridae* and *Rhabdoviridae* families. Importantly, piRNA clusters of *Aedes* mosquitoes are enriched for nrEVE sequences (Aguiar et al., 2020; Crava et al., 2021; Horst et al., 2019; Palatini et al., 2020; Suzuki et al., 2017; Tassetto *et al*., 2019; Whitfield et al., 2017), suggesting that they may serve as the source of immunological memory against exogenous viruses. Indeed, a recent study showed that piRNAs produced from a naturally occurring nrEVE protected the *Ae. aegypti* host against its cognate virus, thus demonstrating a functional link between nrEVEs and antiviral immunity (Suzuki *et al*., 2020).

Given the rapid evolution of the piRNA pathway and the prominent differences between flies and mosquitoes (Assis and Kondrashov, 2009; Wierzbicki et al., 2021), we explored the molecular mechanisms underlying piRNA cluster biogenesis in *Aedes* mosquitoes. We combined transcriptome and chromatin state analyses in *Ae. aegypti* and uncovered readthrough transcription as a major mechanism for piRNA cluster biogenesis. Further comparative analyses between two *Aedes* mosquito species revealed an intriguing model for piRNA cluster evolution in which a conserved set of genes determine the loci for piRNA biogenesis, while piRNA clusters function as traps for nrEVEs, allowing species-specific adaptation to viral infection challenges.

## RESULTS

### Systematic analyses of piRNA clusters in *Ae. aegypti*

As in most arthropods, the piRNA pathway of the vector mosquito *Ae. aegypti* is active in both somatic and germline tissues (Akbari *et al*., 2013; Lewis et al., 2018; Ma *et al*., 2021; Miesen *et al*., 2016b). However, an important question is whether piRNA expression differs across cell lines and tissues in *Ae. aegypti*. To answer this question, we collected piRNA sequencing data from two cell lines (Aag2 (Miesen *et al*., 2016a) and CCL125 (Ma *et al*., 2021)), four somatic tissues (fat body (Zhang et al., 2017), head (Adelman et al., 2013), midgut (Olmo et al., 2018), and thorax (Lewis *et al*., 2018)), as well as two germline tissues (ovary (Lewis *et al*., 2018) and testes), and compared their piRNA cluster expression in a systematic manner (Figure 1A).

**Figure 1.**
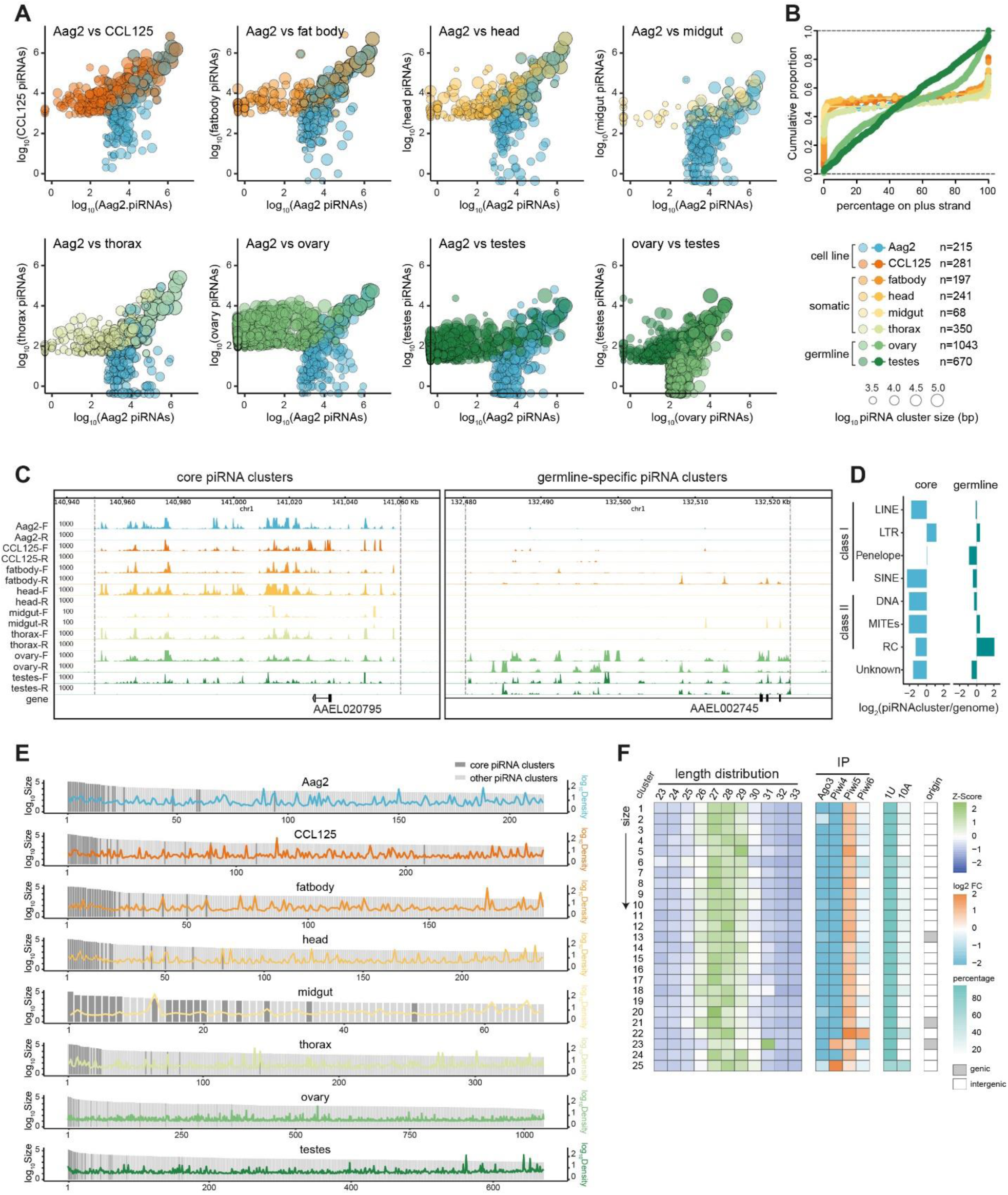
Systematic annotation of piRNA clusters in *Ae. aegypti* mosquitoes. (A) Comparison of piRNA expression from piRNA clusters between cell lines, somatic and germline tissues of *Ae. aegypti*. piRNA clusters were independently annotated in each dataset and represented as dots, with color representing the origin and dot size representing cluster size. The numbers of clusters are listed. Log scaled piRNA read counts were used to represent piRNA expression levels within a piRNA cluster. For detailed information, see Table S1. (B) Cumulative distribution of piRNA cluster coverage on the plus strand of the *Ae. aegypti* genome. Colors are as in (A). (C) Representative genome browser tracks showing examples of a core piRNA cluster and a germline-specific piRNA cluster. piRNA-sized reads (25 to 30 nt) from small RNA-seq were mapped and normalized to RPKM (y-axis), with F/R indicating forward/reverse strand. The boundaries of each cluster are indicated with dashed vertical lines. The boundaries of core piRNA clusters are based on Aag2 piRNA clusters, and germline-specific clusters are based on ovary piRNA clusters. (D) Enrichment of the indicated repeat classes in core and germline-specific piRNA clusters, relative to the entire genome. (E) Distribution of 25 core piRNA clusters in cell lines and tissues. Within each dataset, piRNA clusters are ranked according to log scaled cluster size (bp). The 25 core piRNA clusters are shown as dark grey bars; other piRNA clusters are shown as light grey bars. piRNA density (RPKM) was plotted using the same colors as in (A). (F) piRNA length distribution, PIWI protein association (Halbach *et al*., 2020), 1U/10A bias, and the genomic distribution of the 25 core piRNA clusters, which are ranked according to the cluster size in Aag2 cells.

We first annotated piRNA clusters in each cell line and tissue, using two independent pipelines (Figure S1) (Crava *et al*., 2021; Palatini *et al*., 2020; Rosenkranz and Zischler, 2012). We observed largely comparable piRNA expression between cell lines and somatic tissues, whereas the majority of piRNA clusters in germline tissues showed tissue-dominant expression, which, however, correlated well between ovary and testes (Figures 1A and S1C). Uni-strand clusters were more often observed in cell lines and somatic tissues, whereas dual-strand clusters were predominant in germline tissues (Figure 1B). About 90% of the clusters from cell lines and somatic tissues are uni-strand, whereas 45% and 28% of the clusters are uni-strand in ovary and testes, respectively (Figure S1D). Given these clear differences, we defined two classes of piRNA clusters based on their expression pattern: 25 core piRNA clusters that are universally expressed across cell lines and tissues, and 228 germline-specific piRNA clusters that are uniquely expressed in germline tissues, ovary and testes (Figure 1C).

Unlike piRNA clusters in *D. melanogaster* that are highly enriched for TEs (Brennecke *et al*., 2007), the 25 core piRNA clusters were depleted of TEs except for a minor enrichment of LTR-retrotransposons, while germline-specific piRNA clusters were enriched for Rolling-Circle (RC) transposons when compared to the whole genome (Figure 1D). We also explored the relationship between piRNA cluster size and piRNA density but found no correlation (Figures 1E and S1E). Yet, the 25 core piRNA clusters produced 68% of all cluster-derived piRNAs in Aag2 cells, mainly because they are among the largest piRNA clusters, with the largest cluster spanning ∼180 kb (Figure 1E).

We next characterized the 25 core piRNA clusters in terms of the piRNA length distribution, PIWI-protein association, nucleotide bias, and their genomic distribution (Figure 1F). We found that most mapped small RNAs were 25-30 nt in length and were almost exclusively associated with Piwi5, except for cluster 23 and cluster 25 showing a strong association with Piwi4. Moreover, core cluster-derived piRNAs showed a strong 1U but not 10A bias, suggesting that the core piRNA clusters produce Piwi5-associated primary piRNAs. Remarkably, only three core piRNA clusters are embedded within expressed genes in the same orientation, which raises an interesting question of how the remaining majority of core piRNA clusters that are located in intergenic regions are generated (Figure 1F).

### Comprehensive chromatin profiles of the *Ae. aegypti* genome

The intriguing observation that most core piRNA clusters are located in intergenic regions and give rise to Piwi5-associated primary piRNAs prompted us to test whether these clusters are similar to *flam* (Brennecke *et al*., 2007; Goriaux *et al*., 2014; Malone *et al*., 2009; Prud’Homme et al., 1995). To gain precise insights into transcriptional regulation and chromatin states, we performed ChIP-seq of four histone marks in Aag2 cells, including H3K27ac, associated with active enhancers and promoters; H3K4me3, associated with active promoters; H3K36me3, associated with actively transcribed regions; and H3K9me3, associated with heterochromatin regions (Figure 2A) (Atlasi and Stunnenberg, 2017; Bell et al., 2007; Ernst et al., 2011; Huang et al., 2019b; Narlikar et al., 2013). For all four histone marks, two biological replicates showed strong correlations (Pearson’s r > 0.98), indicating the high reproducibility of these datasets (Figure S2B). Using ChromHMM (Ernst and Kellis, 2017), we annotated five chromatin states based on combinations of four histone marks to define heterochromatin, promoter, enhancer domains, as well as regions associated with active transcription and regions without discernible signals (Figures 2A and S2A).

**Figure 2.**
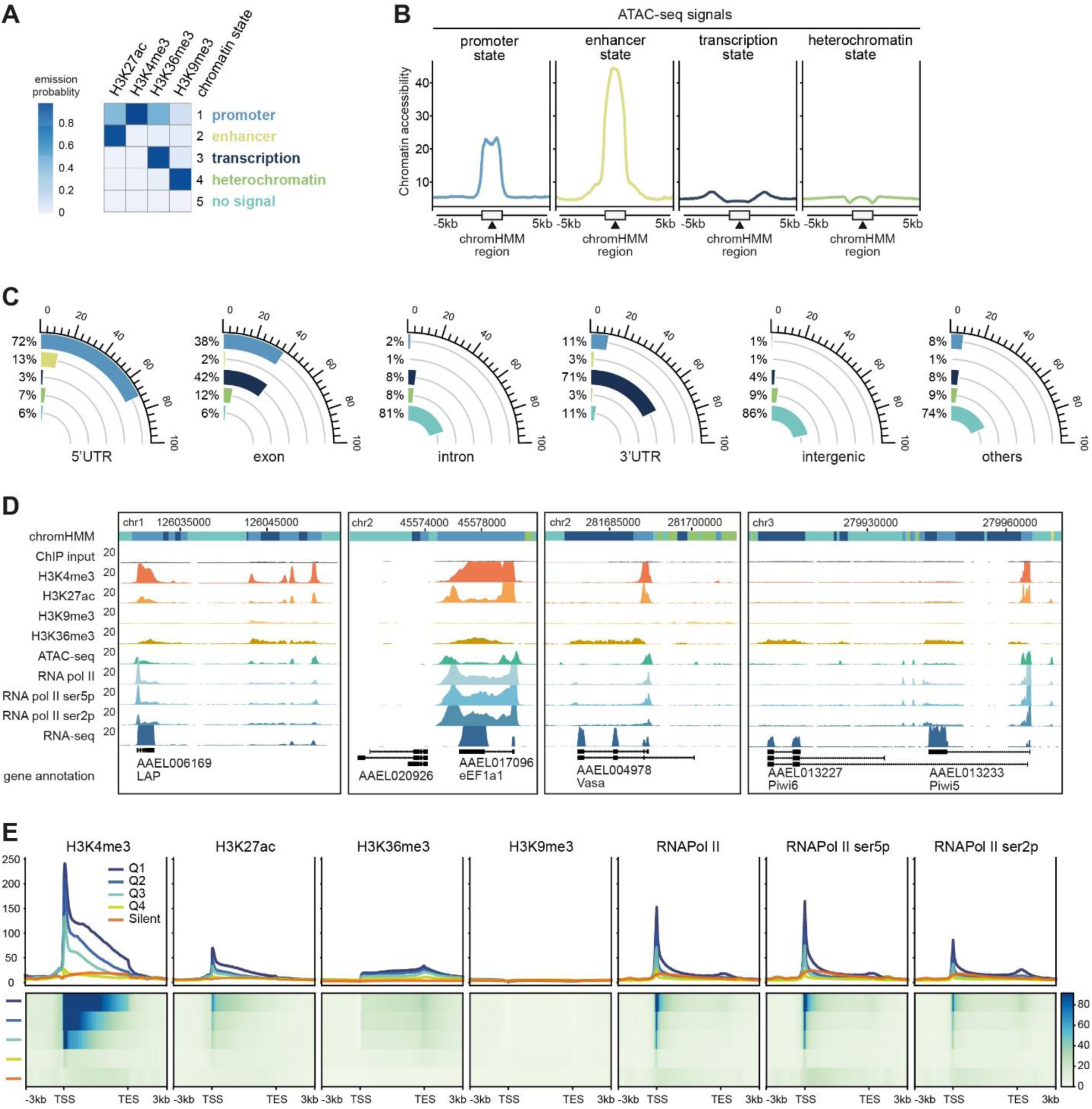
Chromatin profiling in *Ae. aegypti* Aag2 cells. (A) Chromatin state annotation using ChIP-seq data of four histone marks with ChromHMM (Ernst and Kellis, 2017). Each chromatin state (row) is defined by a combinatorial pattern of specific histone marks (columns). For instance, the promoter state is distinguished by enrichment of H3K4me3 while the enhancer state is distinguished by enrichment of H3K27Ac. A five-state ChromHMM was chosen because it represented biologically interpretable patterns that were not repetitive or ambiguous. Six-state and seven-state models are shown in Figure S2A. (B) Average chromatin accessibility marked by ATAC-seq signals at each chromatin state from a. (C) Chromatin state coverage within each genomic feature. When a genomic region was annotated with several chromatin states, only the dominant chromatin state (> 50% coverage of the whole region) is shown. (D) Representative genome browser tracks showing chromatin profiles at expressed genes (*LAP* and *eEF1A1*), a silent gene (*AAEL020926*), genes with different isoforms (*Vasa* and *Piwi6*), as well as genes that are not annotated as such in the reference genome (downstream region of *LAP*). Chromatin state colors are as in a. ChIP-seq and RNA-seq signals were normalized to RPKM (y-axis). (E) Band plots (upper panel) and heatmaps (lower panel) showing average ChIP-seq signals of active histone marks, H3K4me3, H3K27Ac, and H3K36me3, as well as RNA pol II, RNA pol II ser5p, and RNA pol II ser2p. Published RNA-seq datasets were used to define gene expression quartiles (Q1-Q4) and non-expressed genes (silent) (Figure S2E) (Halbach *et al*., 2020). All genes were scaled to 5kb. TSS, transcription start site; TES, transcription end site. ChIP-seq signals were normalized to RPKM (y-axis).

To validate the annotated chromatin states, we performed ATAC-seq to assay open chromatin regions that are biologically active, such as promoters and enhancers (Buenrostro et al., 2013) (Figure 2B). Distinct chromatin accessibility profiles were evident among chromatin states: the promoter and enhancer states showed the highest average levels of chromatin accessibility, followed by regions flanking the transcription state, while the heterochromatin state was relatively inaccessible, in agreement with previous observations in ENCODE datasets (Gorkin et al., 2020). We next assessed the current genome annotation with chromatin states (Figure 2C). In total, 17.4 % of the *Ae. aegypti* genome was assigned with one of the four chromatin states, with most of them located in intron and intergenic regions (Figure S2C). 5’UTRs were enriched for promoter states, while exon and 3’UTR sequences were enriched for the transcription state, and intergenic regions for heterochromatin (Figure 2C).

In addition to histone modifications, eukaryotic transcription units are characterized by conserved 5′ to 3′ profiles of specific RNA polymerase II (pol II) Carboxy-Terminal Domain (CTD) phospho-isoforms (Fong et al., 2017). Phosphorylation of CTD on Ser5 is normally higher at 5′ ends of actively transcribed genes where it facilitates mRNA capping (Schroeder et al., 2000), whereas Ser2 phosphorylation is higher at 3′ ends where it is important for mRNA 3′ end formation (Lunde et al., 2010). We, therefore, performed a series of RNA pol II ChIP-seq experiments, using antibodies either recognizing both phospho-isoforms or one of the two phosphor-isoforms (Figure S2D). We observed that actively expressed genes showed enrichment of active histone marks as well as RNA pol II occupancy at promoter regions, *e.g. LAP* and *eEF1A1*, whereas no signals were found for silenced genes, *e.g. AAEL020926* (Figure 2D). Besides, this dataset provides evidence of differentially expressed isoforms (*Vasa* and *Piwi6*) by marking the corresponding active promoter region. Overall, we observed transcription-dependent signals of active histone marks H3K4me3, H3K27Ac, and H3K36me3, as well as RNA pol II, starting sharp at transcription start sites (TSS) and ending at transcription end sites (TES) (Figure 2E), which has been reported in other organisms (Ando et al., 2019; Lopez-Atalaya et al., 2013; Morselli et al., 2015; Yan et al., 2019). Our genome-wide maps of chromatin accessibility, histone modifications, and RNA pol II occupancy, provide a comprehensive functional annotation of the *Ae. aegypti* genome which serves as a useful resource to understand piRNA cluster biogenesis.

### Readthrough transcription of core piRNA clusters from upstream genes

In contrast to protein-coding genes, we did not detect signals of active histone marks and RNA pol II occupancy at the start of core piRNA clusters (Figure 3A). Instead, active histone marks and RNA pol II signals were detected already in the upstream regions, suggesting that piRNA cluster transcription is initiated from upstream regions and that core piRNA clusters lack their own independent promoters. Besides, we did not detect enrichment of H3K9me3 signals which mark dual-strand clusters in *Drosophila* (Mohn *et al*., 2014), neither for core piRNA clusters nor for other piRNA clusters in Aag2 cells (Figure S3A). Only a few piRNA clusters showed H3K9me3 signals, which, however, coincided with ChIP input signals, and are likely due to the background artifacts from genomic DNA (Amemiya et al., 2019). Together, our analyses revealed that most core piRNA clusters are located in intergenic regions and likely depend on readthrough transcription.

**Figure 3.**
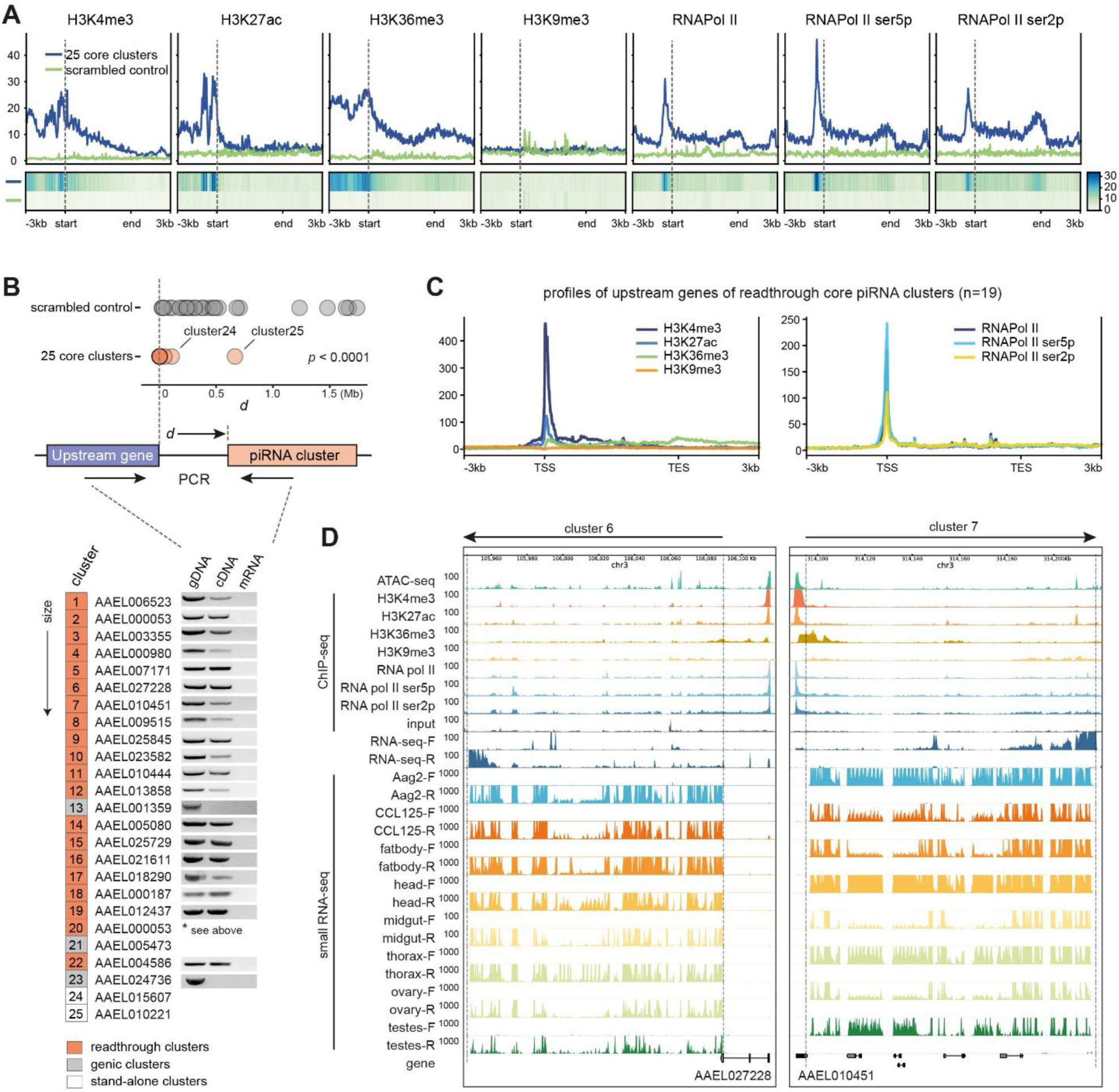
Confirmation of readthrough transcription. (A) Average ChIP-seq signals of active histone marks, H3K4me3, H3K27Ac, and H3K36me3, as well as RNA pol II in 25 core piRNA clusters. All clusters were scaled to 5kb, with dashed lines indicating the 5’ ends of the clusters. Scrambled control regions were generated by shuffling piRNA clusters through the whole *Ae. aegypti* genome with bedtools shuffle (Quinlan and Hall, 2010). (B) The distance (*d*) between the 5’ end of core piRNA clusters to TES of its nearest expressed gene in the same orientation was quantified. Scrambled control regions were generated by shuffling piRNA clusters through the whole genome with bedtools shuffle (Quinlan and Hall, 2010). Statistical differences were examined by unpaired t-test. Lower panel, PCR was performed to analyze readthrough transcription from the upstream gene into the piRNA cluster, using genomic DNA, cDNA, or mRNA as templates. piRNA clusters were ranked according to the cluster size. For detailed primer information, see Table S4. (C) Band plots of average ChIP-seq signals (RPKM) of four histone marks, H3K4me3, H3K27Ac, H3K36me3, H3K9me3 (left), and RNA pol II (right) of 19 upstream genes of readthrough piRNA clusters. All gene body regions were scaled to 5kb. (D) Representative genome browser tracks showing readthrough core piRNA clusters. ATAC-seq, ChIP-seq, and RNA-seq signals, as well as piRNA-sized reads from small RNA-seq, were normalized to RPKM. The boundaries of each cluster are indicated with dashed lines and the direction of piRNA clusters is indicated with arrows, with upstream genes labeled with gene names.

To test this hypothesis, we quantified the distance (*d*) between the 5’ end of core piRNA clusters to the TES of its nearest gene expressed in the same orientation (Figure 3B). Compared with scrambled controls, *d* of 25 core piRNA clusters was significantly shorter. In fact, most core piRNA clusters start directly after the TES of their corresponding upstream genes (*d* = 0), with the exceptions of cluster 24 and cluster 25, which are distant from any expressed genes, which we will refer to as ‘stand-alone’ piRNA clusters (Figure S3B). We noticed that cluster 2 and cluster 20 were mapped to the same upstream gene, *AAEL000053* (Figure 3B). After careful inspection, we propose that these two clusters are part of a large single element, > 220 kb in size, which was annotated as two separate clusters due to a large gap in small RNA coverage (Figure S3B). Moreover, we experimentally validated readthrough transcription using PCR primers spanning the upstream gene and piRNA clusters using cDNA as a template (Figure 3B), whereas properly terminated genes failed to amplify any products (*e.g.*, *AAEL001359* and *AAEL024736*).

We then analyzed chromatin profiles of upstream genes, which showed canonical features of actively transcribed genes, with the enrichment of active histone marks as well as RNA pol II ChIP-seq signals at the TSS (Figure 3C). Expression of several upstream genes, *AAEL006523*, *AAEL005080*, and *AAEL004586*, was confirmed by a recently published proteomic study with > 98% hit identity (Maringer et al., 2017), suggesting these genes are not only transcribed but also translated into proteins. Hence, it is likely that these upstream genes can both deliver properly terminated gene products as well as readthrough transcripts that are substrates for piRNA biogenesis. Notably, most readthrough core piRNA clusters showed higher RNA-seq signals at the end of clusters (Figure 3D), which was also true for genic piRNA clusters (Figure S3B). This observation is likely due to the oligo (dT)-primed RNA-seq library preparation, which in turn suggests that these piRNA cluster transcripts are polyadenylated. Together, our analyses indicate that most core piRNA cluster transcripts are produced by readthrough transcription from upstream genes.

We further dissected the connection between the upstream genes and the downstream piRNA production. Our previous analyses indicated that most core piRNA clusters produce Piwi5-bound primary piRNAs (Figure 1F). We, therefore, used *Piwi5* knockdown as a positive control to establish Stem-Loop (SL)-RT-qPCR assays to quantify the piRNA abundance from readthrough core piRNA clusters (Figure 4A, top panels). Using four piRNAs from two clusters (clusters 7 and 10) as readouts, we analyzed whether silencing of the upstream gene would affect its downstream piRNA production. Upon knockdown of the upstream gene of cluster 7, *AAEL010451*, we did not observe an effect on piRNA expression, probably due to the relatively low knockdown efficiency (Figure 4A, middle panels). We also tried another set of dsRNA targeting *AAEL010451* and did not achieve a better knockdown (Figure S4A). Importantly, upon knockdown of the upstream gene of cluster 10, *AAEL023582*, we observed a significant reduction in piRNA production from its downstream piRNA cluster (Figure 4A, lower panels). This effect was specific, as *AAEL023582* knockdown did not affect the expression of piRNAs from cluster 7. These results provide a direct genetic link between upstream gene transcription and downstream piRNA cluster expression.

**Figure 4.**
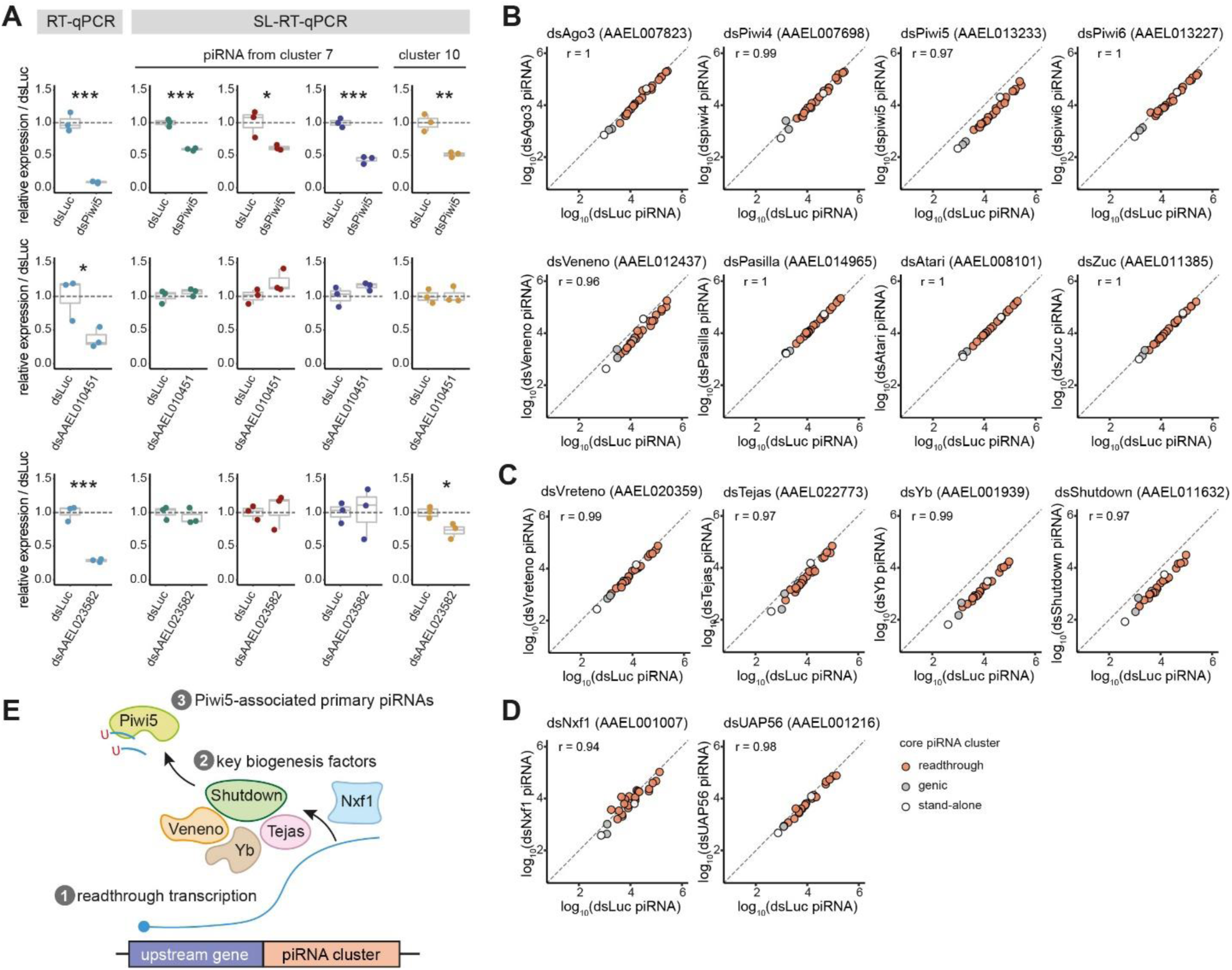
Key determinants of piRNA cluster biogenesis. (A) Knockdown of upstream genes from two readthrough core piRNA clusters. *AAEL010451* is the upstream gene of cluster 7 and *AAEL023582* is the upstream gene of cluster 10. RT-qPCR was performed to confirm gene knockdown, while SL-RT-qPCR was performed to quantify piRNA expression. All experiments were performed in biological triplicates (shown as dots) unless otherwise specified. The interquartile range and the median are shown in box plots. Statistical differences were examined by unpaired t-test, when compared to the non-targeting control knockdown of luciferase (dsLuc). Symbols denote statistical significance (* *p* < 0.05, ** *p* < 0.01, *** *p* < 0.001). For detailed primer information, see Table S4. (B) Changes in piRNA levels from core piRNA clusters upon knockdown of published piRNA biogenesis factors, quantified from small RNA-seq data and compared to the non-targeting control knockdown of luciferase (dsLuc). piRNAs were first normalized to per million mapped miRNAs and then log scaled for comparison. The Pearson correlation coefficient is shown in the upper left corner. (C) Changes in piRNA levels from core piRNA clusters upon knockdown of newly Piwi5 co-factors. Knockdown of target genes was confirmed with RT-qPCR (Figure S4B). (D) Changes in piRNA levels from core piRNA clusters upon knockdown of newly screened piRNA biogenesis factors. Knockdown of target genes was confirmed with RT-qPCR (Figure S4B). (E) A schematic model of piRNA biogenesis from core piRNA clusters in *Ae. aegypti*.

### Key factors of piRNA cluster biogenesis

To define factors involved in the biogenesis of core cluster-derived piRNAs, we analyzed our previously generated small RNA-seq data from Aag2 cell upon knockdown of PIWI genes (Miesen *et al*., 2015), genes encoding PIWI interactors (Joosten *et al*., 2019; Joosten et al., 2021b), *Veneno*, *Pasilla*, *Atari*, as well as *Zucchini* (Zuc), involved in piRNA 3’end formation (Joosten *et al*., 2021a). Among them, we observed a clear reduction in the expression of cluster-derived piRNAs with knockdown of *Piwi5* and *Veneno*, but not with the others (Figure 4B). As most core piRNA clusters are *Piwi5*-dependent (Figure 1F), we further tested three Tudor proteins that were among the top enriched hits from a Piwi5-IP experiment (Joosten *et al*., 2021b), Vreteno (*AAEL020359*), Tejas (*AAEL022773*), and Yb (*AAEL001939*). We also included Shutdown (*AAEL011632*), which is required for the biogenesis of all piRNA populations in *D. melanogaster* (Olivieri et al., 2012). With small RNA-seq, we observed a consistent downregulation of core piRNA clusters upon knockdown of *Yb* and *Shutdown*; in comparison, the effect of Tejas was minor and Vreteno barely affected piRNA expression from these core piRNA clusters (Figure 4C).

To search for key factors that are upstream in the piRNA pathway, we performed a large-scale RNAi screen, including orthologs of known piRNA biogenesis factors in other organisms, chromatin modifiers, 16 genes of mRNA 3’end processing, 18 genes that mediate RNA export, 26 genes of general RNA processing, as well as 29 RNA helicases (Figure S4). With the screen, we found that *Nxf1* knockdown significantly downregulated one of the two tested piRNAs (Figure S4E). We then confirmed the effect of *Nxf1* knockdown using another set of dsRNA and observed a significant reduction of two piRNAs from two piRNA clusters. Next, we performed small RNA-seq for a more comprehensive evaluation of all core piRNA clusters (Figure 4D). In addition, we included UAP56 which plays an important role in piRNA cluster export in *D. melanogaster* (ElMaghraby *et al*., 2019). The effect of *Nxf1* knockdown seemed to be more heterogeneous, however, not correlated to whether the cluster is readthrough, genic or stand-alone; whereas, knockdown of *UAP56* barely affected piRNA expression from these core piRNA clusters, in line with results from the SL-RT-qPCR screen (Figure S4E).

Together, these data suggest the following model for cluster-derived piRNA biogenesis: 1) piRNA cluster transcripts are produced by readthrough transcription from an upstream coding gene; 2) key factors involved in processing and maturation include the mRNA exporter Nxf1, three Tudor proteins, Veneno, Tejas, and Yb, the evolutionarily conserved cochaperone Shutdown, as well as the PIWI protein Piwi5; 3) the resulting mature piRNAs are mainly associated Piwi5 and show a strong 1U bias (Figure 4E).

### Conserved readthrough transcription for piRNA cluster biogenesis among mosquitoes

To analyze whether readthrough transcription is a shared mechanism for cluster piRNA biogenesis, we extended our analyses to other mosquito species, including the closely related *Aedes albopictus*, and the more distant *Culex quinquefasciatus* and *Anopheles gambiae*, (Figure 5A). With the same strategy used for *Ae. aegypti* (Figure 1), we annotated piRNA clusters from cell lines and tissues from these mosquito species (Figures S5-7). Like in *Ae. aegypti*, most piRNA clusters were uni-strand in cell lines and somatic tissues, whereas in germline tissues piRNA clusters are more dual-stranded. About 70% of the clusters from cell lines and somatic tissues were uni-strand, whereas the percentage was below 52% for germline tissues in all three species. Next, we defined core piRNA clusters in *Ae. albopictus* (n=37), in *Cu. quinquefasciatus* (n=32), and in *An. gambiae* (n=19), based on their ubiquitous expression across different cell lines and tissues. In parallel, we defined germline-specific piRNA clusters in *Ae. albopictus* (n=244), in *Cu. quinquefasciatus* (n=587), and in *An. gambiae* (n=539) (Tables S5-7).

**Figure 5.**
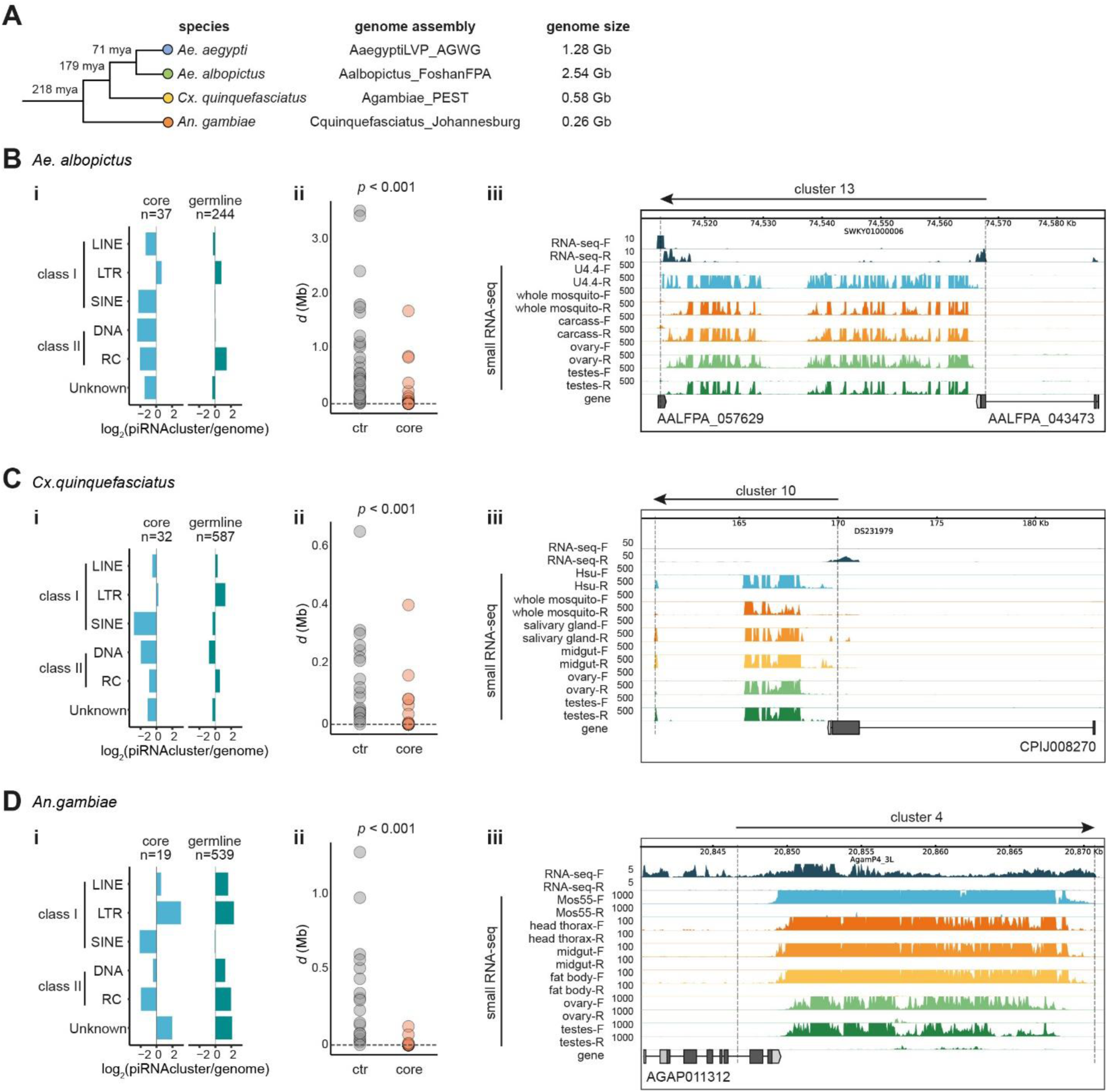
Readthrough transcription is an evolutionarily conserved mechanism for piRNA cluster biogenesis in mosquitoes. (A) Phylogeny and divergence time estimation by molecular clock analysis of four mosquito genomes (Chen *et al*., 2015), with the genome assembly (Vectorbase release 56) and the corresponding genome size annotated. (B)-(D) Analyses of piRNA clusters in *Ae. albopictus* (B), *Cx. quinquefasciatus* (C), and *An. gambiae* (D), including enrichment of repeat classes in piRNA clusters (i), the distance (*d*) as illustrated in Figure 3B (ii), and representative readthrough core piRNA clusters (iii).

Analogous to *Ae. aegypti*, core piRNA clusters in these three species were only modestly enriched for LTR retrotransposons (Figures 5B-D), whereas germline-specific piRNA clusters in *Ae. albopictus* and *Cx. quinquefasciatus* mainly showed slight enrichment of LTR retrotransposons and RC transposons (Figure 5B and c). In striking contrast, piRNA clusters in *An. gambiae* showed an overall enrichment of all TE classes, except for SINEs (Figure 5D). Core piRNA clusters in *Ae. albopictus* were also among the top-sized piRNA clusters (Figure S5D), whereas core piRNA clusters in *Cx. quinquefasciatus* and *An. gambiae* were more dispersed in size (Figures S6D and S7D).

We next examined whether core piRNA clusters follow the pattern of readthrough transcription, using *d* (illustrated in Figure 3B) as an indicator. Compared to scrambled controls, *d* of core piRNA clusters was significantly shorter in all three species (Figures 5B-D). To define whether a core piRNA cluster follows readthrough transcription, we set a cutoff of *d* based on the *d* in *Ae. aegypti* (5 kb), scaled by the genome size of each mosquito species (Figure 5A) to account for the higher probability to classify a piRNA cluster as a readthrough cluster in smaller genomes (*d* < 10 kb for *Ae. albopictus*; 2.5 kb for *Cx. quinquefasciatus*, and 1.0 kb for *An. gambiae*). In summary, 14 of 37 core piRNA clusters in *Ae. albopictus* (Figure S5E), 14 of 32 core piRNA clusters in *Cx. quinquefasciatus* (Figure S6E), and 10 of 19 core piRNA clusters in *An. gambiae* (Figure S7E), fulfilled the criteria and are thus likely transcribed by readthrough transcription from upstream genes. Indeed, many of these core piRNA clusters started directly after the upstream genes (Figures 5B-D). In conclusion, these analyses suggest that readthrough transcription is a common mechanism for piRNA cluster biogenesis shared among these four vector mosquitoes.

### A conserved pool of genes drives piRNA cluster expression in *Aedes* mosquitoes

Our observation that readthrough transcription is a conserved mechanism for piRNA biogenesis raises the question of whether specific genes are selected to drive cluster expression. For the two *Aedes* mosquitoes, the expression of upstream genes was significantly lower when compared to all expressed genes, whereas the gene size seemed to be larger, however, the difference was not significant (Figure 6A). In contrast, the upstream genes of *Cx. quinquefasciatus* and *An. gambiae* were expressed at higher levels. With Gene Ontology (GO) analysis, the upstream genes in *Cx. quinquefasciatus* were significantly enriched for genes encoding ribosomal proteins, whereas no GO term enrichment was found in any other species.

**Figure 6.**
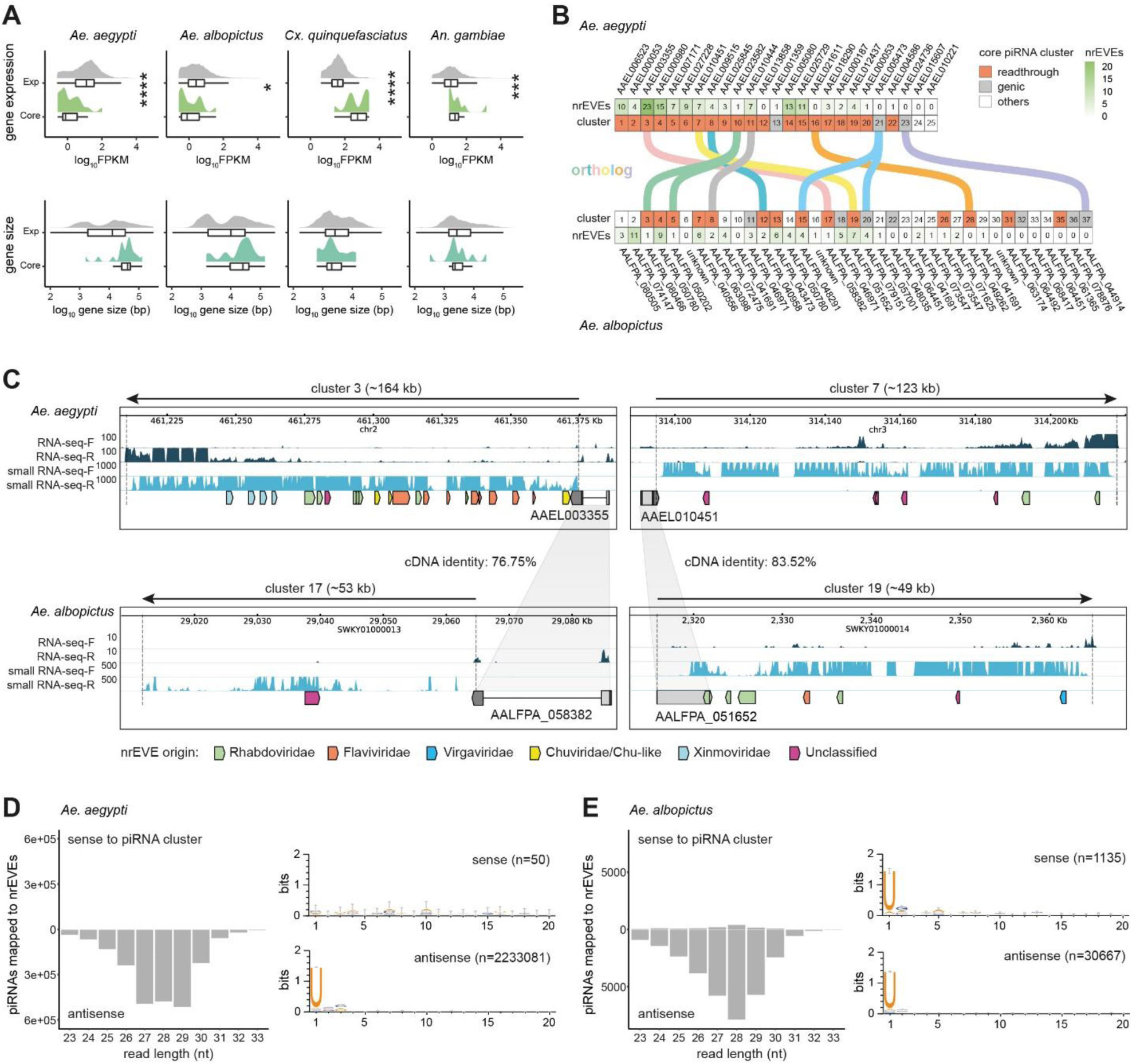
Systematic comparison of core piRNA clusters between mosquitoes. (A) Expression level and the gene body length of upstream genes of core piRNA clusters (Core) compared to all expressed genes (Exp) in each species. Expressed genes were determined for each species (Figure S8A). Both the distribution of the raw data and the box plot displaying the interquartile range and the median are shown. Statistical differences were examined by unpaired t-test (* *p* < 0.05, *** *p* < 0.001, **** *p* < 0.0001). (B) Comparison between core piRNA clusters in two *Aedes* mosquitoes. In each species, piRNA clusters were ranked according to their size and colored based on their genomic origin, with another column showing the number of nrEVE insertions. Ortholog genes were connected. ‘Unknown’ refers to the situation when a piRNA cluster is located in a scaffold without any annotated gene nearby. (C) Representative genome browser tracks showing two pairs of core piRNA clusters with the upstream genes being orthologs in two *Aedes* mosquitoes. The boundaries of each cluster are indicated with dashed vertical lines and the direction of the piRNA cluster is indicated with an arrow on top. nrEVEs were shown as arrows, with colors representing viral families. (D) Read size distribution and the nucleotide bias of piRNAs mapped to nrEVEs within 25 core piRNA clusters of *Ae. aegypti*. (E) Read size distribution and the nucleotide bias of piRNAs mapped to nrEVEs within 37 core piRNA clusters of *Ae. albopictus*.

Notably, many upstream genes are orthologs between *Ae. aegypti* and *Ae. albopictus*, which is also true for a few genic piRNA clusters (Figure 6B). This allowed us to define homologous piRNA clusters and compare their sequence composition in a pair-wise manner (Figure 6C). Despite the highly conserved upstream genes (cDNA identity > 75%), the downstream piRNA clusters differed both in size and sequence identity. The latter was especially evident with varied numbers of nrEVE insertions from different virus families into orthologous clusters (Figures 6B and C, summarized in Table S8). Gene duplication is a common phenomenon in the genome of *Ae. albopictus* Foshan strain (Chen et al., 2015) and we observed an interesting example where in *Ae. aegypti*, *AAEL005473* was associated with a genic piRNA cluster (cluster 21) and depleted of nrEVE insertions, whereas its two orthologous genes in *Ae. albopictus* were associated with a readthrough piRNA cluster (cluster 15) and a genic piRNA cluster (cluster 20), both with nrEVE insertions (Figure S8B).

Several studies have shown that piRNA clusters are enriched for nrEVEs in *Aedes* mosquitoes (Crava *et al*., 2021; Horst *et al*., 2019; Palatini *et al*., 2020; Palatini et al., 2017; Whitfield *et al*., 2017). We revisited the annotated nrEVEs in *Ae. aegypti* (Crava *et al*., 2021) and found that 133 of 252 nrEVEs (53%) are located within 215 piRNA clusters of Aag2 cells, and 126 of these 133 nrEVEs (95%) are located within the 25 core piRNA clusters (*X^2^* test, *p* < 0.0001) (Figure 6B). The same is true for *Ae. albopictus* (Palatini *et al*., 2020), in which 119 of 456 nrEVEs (26%) are located within 285 piRNA clusters, and among them, 82 of these 119 nrEVEs (69%) are located within the 37 core piRNA clusters (*X^2^* test, *p* < 0.0001). Besides, the vast majority of nrEVEs were inserted in antisense orientation to core piRNA clusters in both species and thus serve as a pool of primary piRNAs that can potentially target cognate virus sequences (Figures 6D and E). In *Ae. albopictus*, there are two exceptions where nrEVEs insertions were in the same orientation as core piRNA clusters, such as the nrEVE derived from a member of the *Rhabdoviridae* within cluster 20 (Figure S8B), the resulting piRNAs were sense to piRNA clusters, yet also showed a strong 1U bias (Figure 6E).

Together, these analyses revealed that a conserved pool of genes was selected to define the genomic loci for piRNA cluster biogenesis in *Aedes* mosquitoes; however, the sequence composition of piRNA clusters is not conserved and core clusters have evolved as ‘traps’ for nrEVE insertions in a species-specific manner. Although the mechanism of readthrough transcription is conserved, the identity of upstream genes is not conserved in evolutionary more distant mosquito species.

### Core piRNA clusters function as traps for nrEVEs

We further explored why nrEVEs are preferentially inserted into core piRNA clusters in *Aedes* mosquitoes. As nrEVE insertions and piRNA cluster evolution are highly intertwined processes in both *Aedes* mosquitoes, we included *Cx. quinquefasciatus* and *An. gambiae* in this analysis, because their genomes contain few nrEVE insertions and, therefore, their core piRNA clusters likely represent the naïve state of core piRNA clusters devoid of nrEVE insertions (Palatini *et al*., 2017). We compared core piRNA clusters with the remaining piRNA clusters in terms of cluster size and piRNA density. We found core piRNA clusters are larger in cell lines and tissues of all four species, except for two germline tissues of *An. gambiae* (Figure 7A). The differences in cluster size are more drastic in the two *Aedes* mosquitoes than in *Cx. quinquefasciatus* and *An. gambiae*. We observed a general pattern of higher piRNA density in core piRNA clusters of all four species. Thus, a possible scenario is that core piRNA clusters are intrinsically larger in size and high in piRNA density which favors insertions of nrEVEs; as a consequence, nrEVE insertions enlarge core piRNA clusters without adversely affecting piRNA production and drive core piRNA cluster expansion.

**Figure 7.**
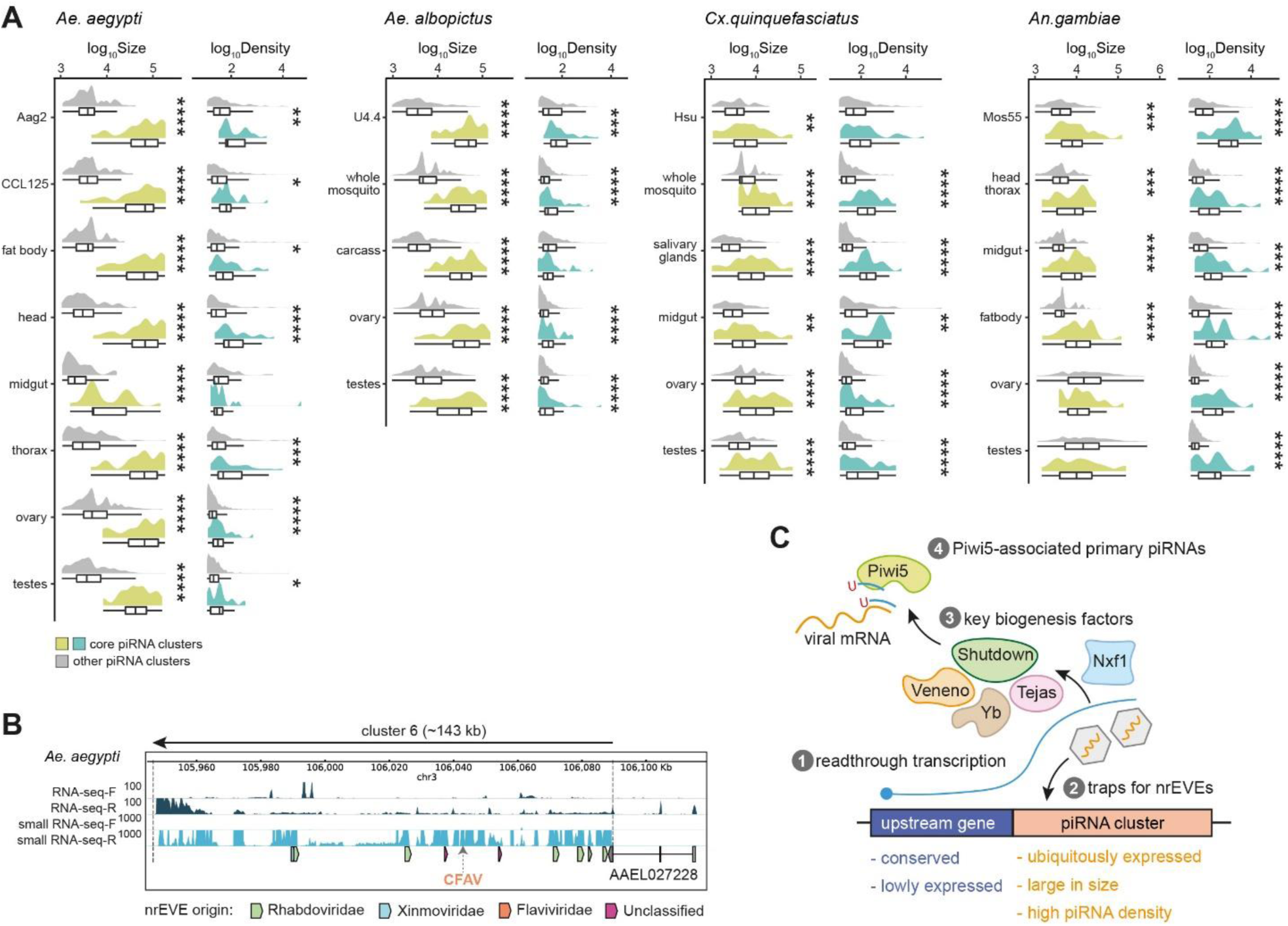
Core piRNA clusters as traps for nrEVEs. (A) Comparison of log scaled piRNA cluster size (bp) and piRNA density (RPKM) between core piRNA clusters and the remaining piRNA clusters across different cell lines and tissues in four species. Statistical differences were examined by unpaired t-test (* *p* < 0.05; ** *p* < 0.01; *** *p* < 0.001; **** *p* < 0.0001). (B) Representative genome browser tracks showing a CFAV-derived nrEVE insertion into the core piRNA cluster 6 of *Ae*. *aegypti* from wild-caught mosquitoes (Crava *et al*., 2021). The boundaries of the piRNA cluster are indicated with dashed lines and the direction of the piRNA cluster is indicated with an arrow on top. nrEVEs are shown as arrows, with colors representing viral families. (D) Schematic model for piRNA cluster biogenesis in *Aedes* mosquitoes.

The acquisition and loss of nrEVEs from the host genome are highly dynamic and ongoing processes. Recent studies identified nrEVEs that were absent from the reference genome but present in wild-collected mosquitoes from different geographical populations (Crava *et al*., 2021; Suzuki *et al*., 2020). Among these newly identified nrEVEs, three nrEVEs have definite chromosomal integration sites, one of which was previously been shown to be in a piRNA cluster (Crava *et al*., 2021), which is one of the core piRNA clusters (cluster 6, Figure 7B). With the limited number of defined integration sites for new nrEVEs, it is impossible to perform statistics, but it is worth noting that the 25 core piRNA clusters in total span ∼1.9 Mb, occupying only 0.15% of the *Ae. aegypti* genome. We, therefore, propose that the insertion and maintenance nrEVEs in core piRNA clusters are unlikely to be stochastic, but rather a result of positive selection.

## DISCUSSION

Here, we comprehensively analyzed piRNA expression in four mosquito species and uncovered readthrough transcription as a conserved mechanism for cluster piRNA biogenesis (Figure 7C). Within *Aedes* mosquitoes, the upstream genes are relatively lowly expressed, but evolutionary conserved to drive piRNA cluster transcription. The downstream piRNA clusters are unidirectional, ubiquitously expressed, and function as traps for nrEVEs in a species-specific manner. The piRNA cluster transcripts are processed by key factors, including the RNA export factor Nxf1, three Tudor proteins, Veneno, Tejas, and Yb, as well as the cochaperone Shutdown. Eventually, mature piRNAs antisense to nrEVEs are loaded into Piwi5 and hold the potential to target cognate viral mRNA. Our study proposes a model for piRNA cluster biogenesis in important vector mosquitoes and provides insights into the adaptive evolution of mosquito genomes via the endogenization of viral sequences.

### Characterization of mosquito piRNA clusters

We defined two different categories of piRNA clusters: core piRNA clusters that are shared between somatic and germline tissues; and germline-specific piRNA clusters that show restricted expression only in germline tissues. Core piRNA clusters are predominantly uni-strand clusters and produce Piwi5-bound primary piRNAs independent of the ping-pong amplification loop, thus resembling the *flam* locus in *D. melanogaster* (Brennecke *et al*., 2007; Malone *et al*., 2009). However, there are essential differences. First, *flam* serves as a piRNA cluster only in somatic follicular cells of *Drosophila* ovaries (Brennecke *et al*., 2007), while in somatic tissues outside of the ovary *flam* acts as a source of endogenous siRNAs (Ghildiyal et al., 2008). In contrast, core piRNA clusters in mosquitoes show ubiquitous expression in both somatic and germline tissues. We cannot exclude that the observed expression of core piRNA clusters in germline tissues might result from the residential somatic cell populations, rather than from the actual germline cells, which deserves further investigation. Second, unlike *flam*, which is transcribed by RNA pol II from its own promoter similar to protein-coding genes (Goriaux *et al*., 2014), core piRNA clusters do not have their own promoters, but depend on readthrough transcription from their upstream genes (Figure 3). Third, *flam* is enriched for *gypsy* family LTR retrotransposons, whereas core piRNA clusters are enriched for nrEVEs in *Aedes* mosquitoes (Figure 6) (Crava *et al*., 2021; Palatini *et al*., 2020). Thus, *flam* represses transposons by preventing the expression of *gypsy* family TEs having complementary sequences in this locus, while core piRNA clusters with nrEVEs trapped in the antisense orientation encode a pool of piRNAs that have the potential to target their cognate viruses. A recent study provided direct evidence of nrEVEs mediated antiviral immunity by showing that a naturally occurring nrEVE produces piRNAs to protect the host against its cognate virus, the cell-fusing agent virus (Suzuki *et al*., 2020). Third, the enrichment of the *gypsy* TE family is conserved in *flam* in three closely related species, *D. melanogaster*, *D*. *yakuba*, and *D*. *erecta* (Malone *et al*., 2009). Moreover, *flam* acquires new TEs via horizontal transfer, with related elements showing higher sequence identity between *D. simulans*, *D. sechellia*, *D. yakuba*, and *D. erecta* than expected if TEs would be acquired through vertical transmission (Malone *et al*., 2009; Zanni et al., 2013). In contrast, nrEVE insertions vary both in numbers and virus families between *Ae. aegypti* and *Ae. albopictus* (Figure 6). Therefore, it is likely that core piRNA clusters acquire new nrEVEs via direct endogenization of viral sequences, showing adaption to local viral challenges.

Germline-specific piRNA clusters produce piRNAs from both genomic strands and show restricted expression only in germline tissues. Hence, it is logical to hypothesize that germline-specific piRNA clusters in mosquitoes may resemble dual-strand piRNA clusters expressed in the ovary of *D. melanogaster* that are marked by heterochromatic H3K9me3 (Andersen *et al*., 2017; Klattenhoff *et al*., 2009; Mohn *et al*., 2014; Zhang *et al*., 2014). However, no orthologs have been found in mosquitoes for those key factors that promote dual-stranded piRNA cluster transcription, such as the RDC complex (Chen *et al*., 2016) and Moonshiner (Andersen *et al*., 2017). Besides, there is a clear depletion of most TEs in germline-specific piRNA clusters in mosquitoes with the exception of *An. gambiae* (Figure 5D). Therefore, chromatin profiles in germline tissues are needed to study the regulation of germline-specific piRNA clusters. Recent studies in *D. melanogaster* male germline revealed a sex-specific piRNA program (Chen et al., 2021a; Chen et al., 2021b), which is different from our observations in mosquitoes where piRNA clusters correlated well between ovaries and testes, which was especially obvious in *An. gambiae* (Figure S7A). Future studies of germline-specific piRNA clusters are required to gain more insights into the involved key factors and the underlying mechanism of cluster transcription.

In addition to core piRNA clusters and germline-specific piRNA clusters, we observed an interesting tissue-specific expression pattern that fewer piRNA clusters are expressed in the midgut. Such restricted piRNA expression in the midgut from our analysis is supported by a recent study of dengue virus-infected *Ae. albopictus*, in which it was shown that piRNAs were reduced in the midgut (Wang et al., 2018). Interestingly, another recent study showed that the siRNA pathway fails to efficiently silence dengue virus replication in the midgut of *Ae. aegypti* (Olmo *et al*., 2018). As the midgut is one of the main tissue barriers for arboviruses to cross to establish systematic infection (Franz et al., 2015), it will be interesting to investigate the interplay between these two small RNA pathways in the midgut and to address their role in antiviral immunity.

### Readthrough transcription as a major mechanism for piRNA biogenesis in mosquitoes

Through combined transcriptome and chromatin state analyses of the *Ae. aegypti* genome, we uncovered readthrough transcription as a major mechanism for piRNA cluster biogenesis. Knockdown of one of the upstream genes reduced the corresponding downstream piRNA production. Moreover, many upstream genes are actually orthologs between the two *Aedes* mosquitoes; therefore, readthrough transcription from this subset of genes is definitely not a stochastic event, but rather a tightly regulated and evolutionarily conserved process.

Two connected questions remain to be answered: how these upstream genes are regulated to allow readthrough transcription and how are readthrough transcripts funneled into the piRNA pathway. In germline cells of *D. melanogaster*, the dual-stranded piRNA cluster *hsp70* locus has been proposed to follow readthrough transcription, however, a specialized RDC complex is engaged to inhibit termination by recruiting the cleavage and polyadenylation specificity factor (CPSF) complex in a chromatin-dependent but sequence-independent manner (Chen *et al*., 2016; Mohn *et al*., 2014). We screened 16 genes that are involved in canonical mRNA 3’end processing including all key components of the CPSF complex and did not observe a clear effect. Moreover, core piRNA clusters in mosquitoes are not marked by any repressive histone mark, thus devoid of recognizable chromatin features. Therefore, it is likely that a specialized non-canonical 3’end processing pathway is recruited in a sequence-dependent manner for readthrough transcription of piRNA cluster transcripts.

After being transcribed, the *flam* piRNA precursor is spliced, capped, polyadenylated, and exported by Nxf1/Nxt1 to Yb-bodies where primary piRNA biogenesis occurs (Dennis *et al*., 2016; Goriaux *et al*., 2014; Li *et al*., 2009a; Qi *et al*., 2011). We screened 18 genes of the RNA export pathway and identified Nxf1, but not Nxt1, as an important factor for piRNA clusters. This is rather unexpected because the heterodimeric mRNA export receptor Nxf1-Nxt1 is conserved from budding yeast to humans to mediate mRNA translocation through nuclear pore complexes into the cytoplasm (Katahira et al., 1999). Instead, this may indicate that a specialized non-canonical mRNA export pathway is employed to convey these readthrough transcripts to the piRNA pathway, which involves Nxf1 but not its canonical co-factors, such as Nxt1. So far, it is not clear at which stage piRNA cluster precursors are funneled into the piRNA biogenesis pathway. It is possible that, directly after readthrough transcription, piRNA cluster precursors are uncoupled from the upstream gene transcript and are separately exported by Nxf1. Another possibility is that readthrough transcripts are exported as a whole, and afterwards, the piRNA cluster precursors are recognized by the piRNA pathway for further processing.

Our analyses identified three Tudor proteins, Veneno, Tejas, Yb, and an evolutionarily conserved cochaperone Shutdown, as key factors for piRNA cluster biogenesis. Among them, Tejas and Yb are Piwi5 interactors, Veneno and Shutdown are Ago3 interactors, whereas Veneno also interacts with Yb which in turn interacts with Piwi5 (Joosten *et al*., 2019; Joosten *et al*., 2021b). Such a Yb-centered protein network suggests that Yb bodies are the place for primary piRNA biogenesis in mosquitoes. However, Vreteno, another Tudor protein, did not affect piRNA production, although it is an important component of Yb bodies in *D. melanogaster* (Handler et al., 2011). Therefore, further characterization is needed to dissect the protein composition of Yb bodies in *Aedes* mosquitoes.

### Core piRNA clusters function as traps for nrEVEs

Our analysis revealed that core piRNA clusters function as traps for nrEVEs in the two *Aedes* mosquitoes and that they are evolving in a species-specific manner. The different numbers and identities of acquired nrEVEs likely reflect their exposure to virus challenges, which is supported by the finding of wild-caught mosquito populations from epidemic regions showing a CFAV-derived nrEVE trapped in the core piRNA cluster 6 (Crava *et al*., 2021). Importantly, a recent study provided direct evidence of nrEVEs mediated antiviral immunity by showing that a naturally occurring CFAV-derived nrEVE produces piRNAs to protect the host against its cognate virus (Suzuki *et al*., 2020). Thus, nrEVEs that have been trapped in core piRNA clusters can be regarded as a sequence-specific record of virus exposure that may function in concert with the siRNA machinery which is a broadly active antiviral response.

Core piRNA clusters are intrinsically larger in size and high in piRNA density which, somehow, favors insertions of nrEVEs. Such direct integration, transcription, and production of nrEVE-derived piRNAs constitutes a mechanism of adaptive genome evolution of *Aedes* mosquitoes. Therefore, one possibility is to artificially mimic this heritable immune memory, for example, by inserting sequences of pathogenic arboviruses such as Zika and dengue virus into core piRNA clusters to generate mosquitoes resistant to infection as an intervention strategy.

Our study provides a systematic evaluation of piRNA profiles in four mosquito species and uncovered a conserved readthrough transcription mechanism for piRNA biogenesis. We propose a model for piRNA cluster evolution and highlight the function of piRNA clusters as traps for nrEVEs to shape the antiviral immunity of *Aedes* mosquitoes. Our study lays a solid foundation for future piRNA studies to understand how the piRNA pathway affects the vector competence of mosquito populations via nrEVE insertions, and how nrEVE-derived piRNAs influence the evolution of viruses. Moreover, our genome-wide maps of chromatin state, genome accessibility, and pol II occupancies will be a useful resource for understanding gene regulation in the important vector mosquito *Ae. aegypti*.

## STAR METHODS

### Key resources table

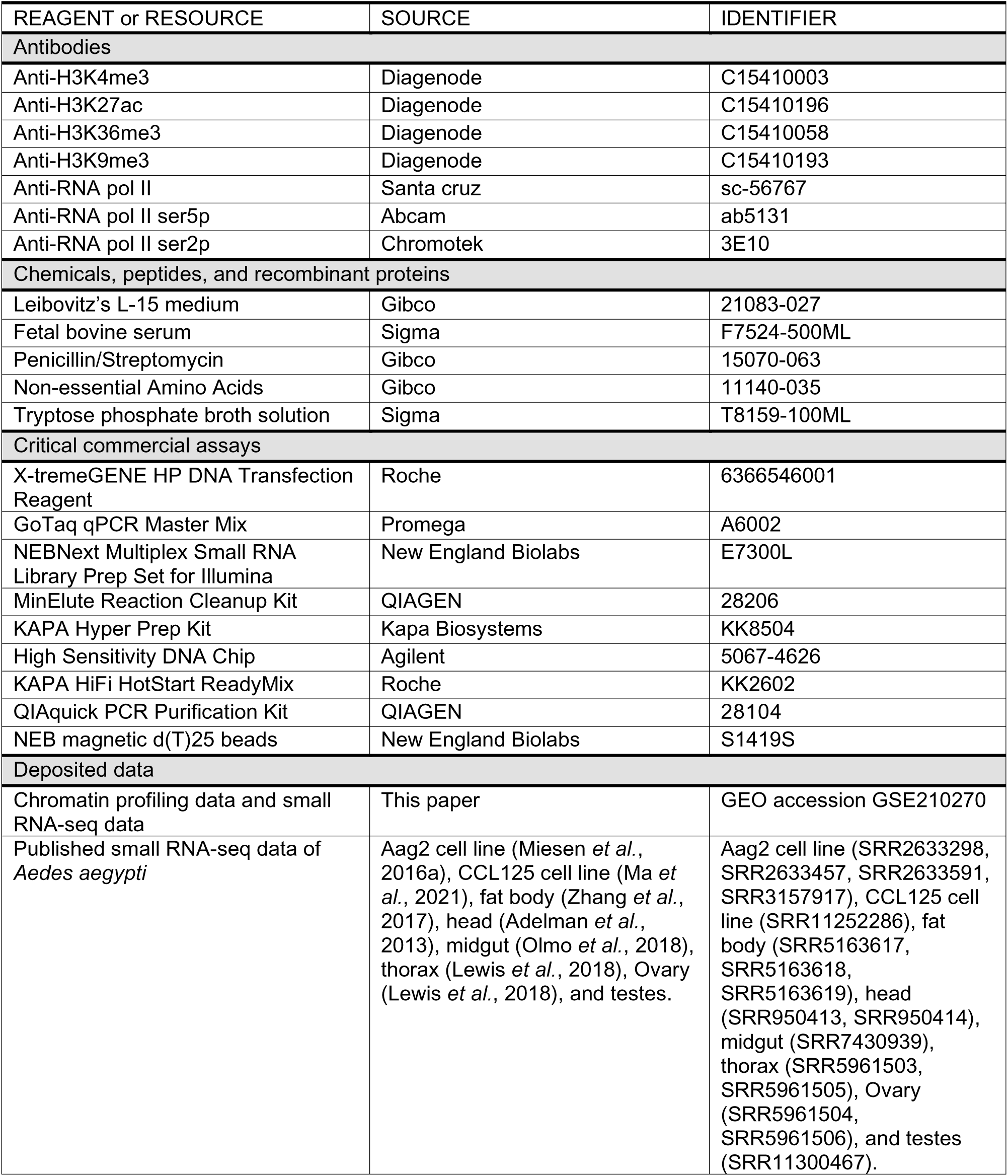
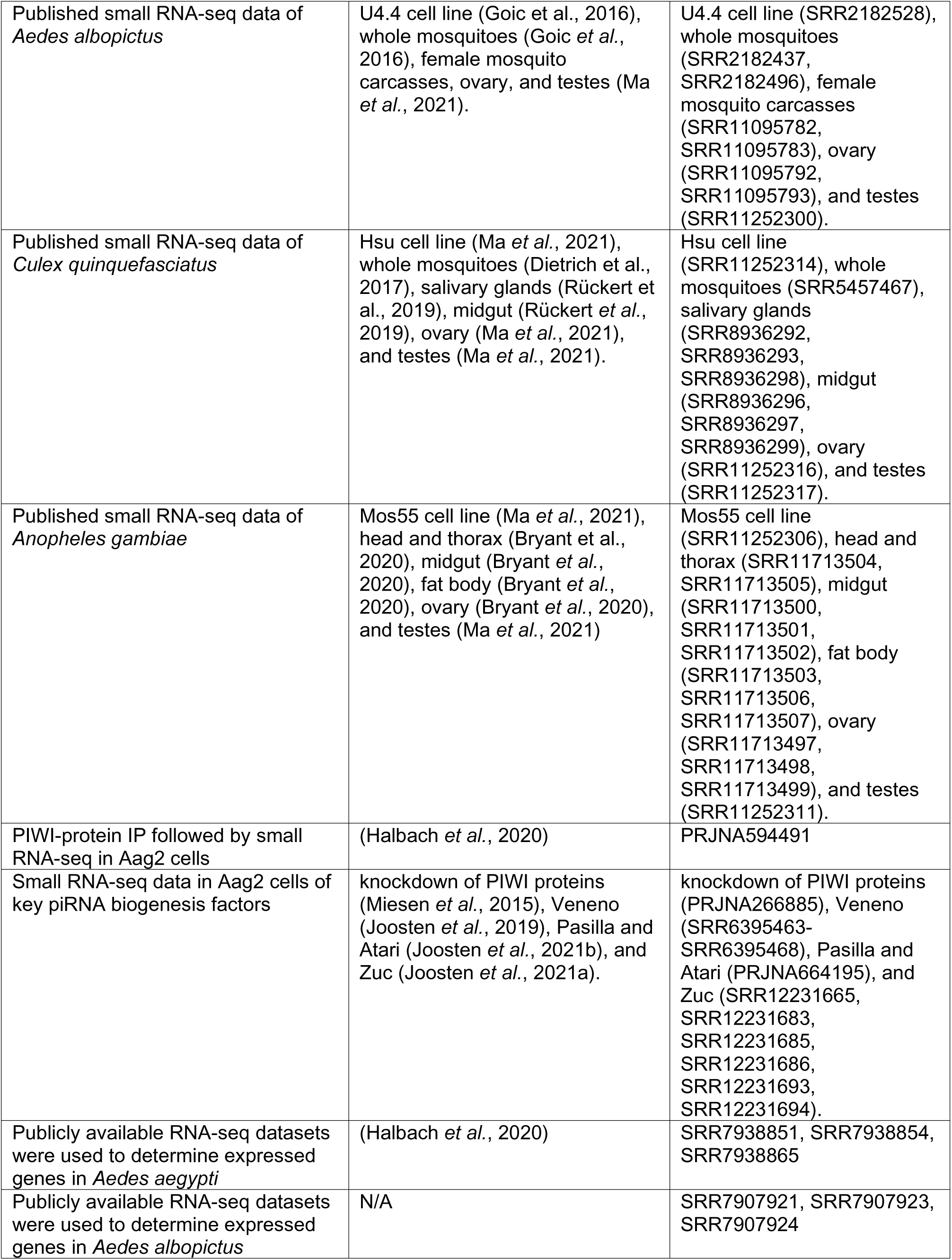
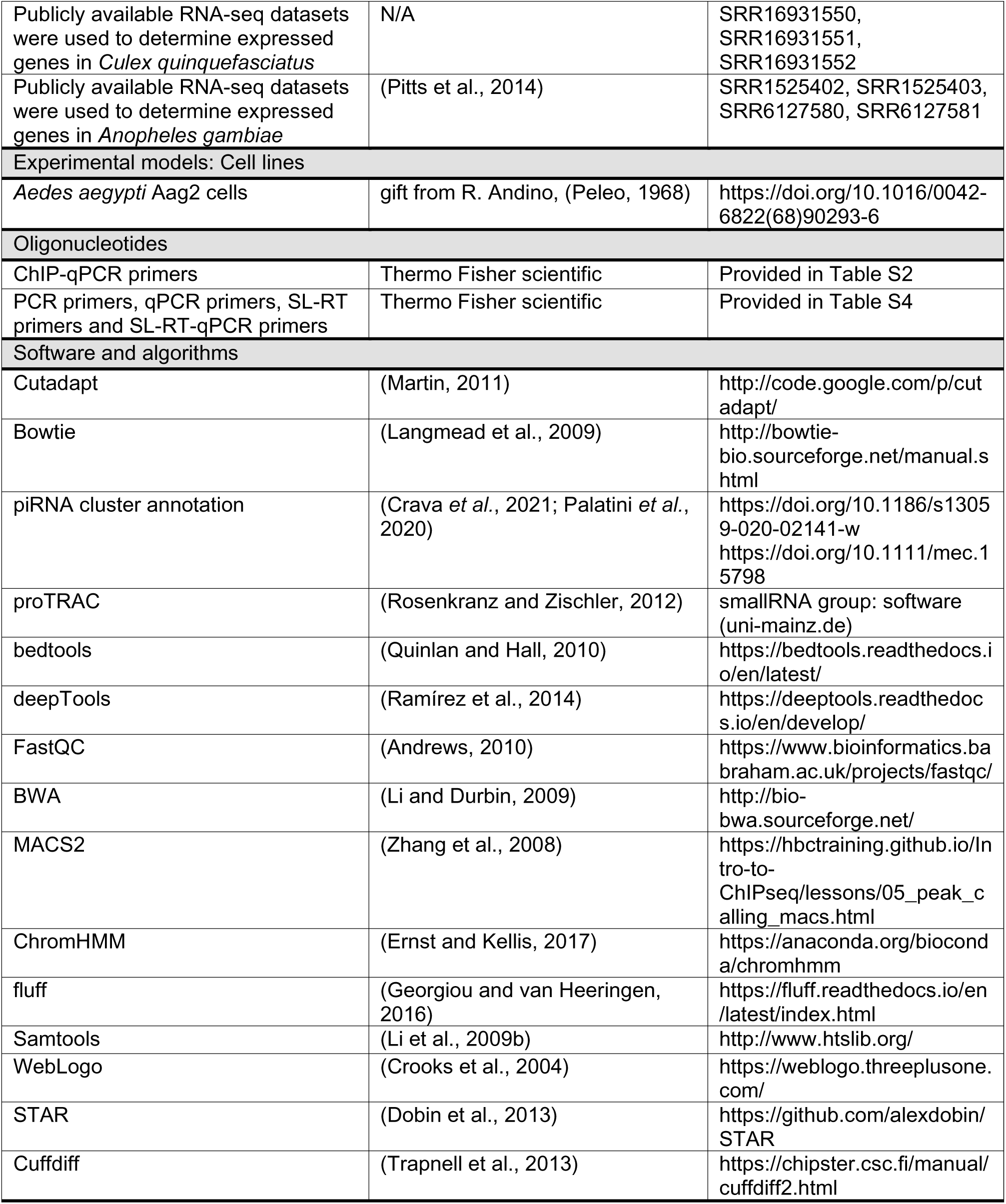

### Method details

#### piRNA cluster annotation

The genome assemblies of four mosquito species, *Ae. aegypti* LVP, *Ae. albopictus* Foshan FPA, *Cx. quinquefasciatus* Johannesburg, *An. gambiae* PEST, were downloaded from Vectorbase (release 56). Publicly available small RNA-seq datasets were collected for piRNA cluster annotation. For *Ae. aegypti*, Aag2 cell line (SRR2633298, SRR2633457, SRR2633591, SRR3157917) (Miesen *et al*., 2016a), CCL125 cell line (SRR11252286) (Ma *et al*., 2021), fat body (SRR5163617, SRR5163618, SRR5163619) (Zhang *et al*., 2017), head (SRR950413, SRR950414) (Adelman *et al*., 2013), midgut (SRR7430939) (Olmo *et al*., 2018), thorax (SRR5961503, SRR5961505) (Lewis *et al*., 2018), Ovary (SRR5961504, SRR5961506) (Lewis *et al*., 2018), and testes (SRR11300467). For *Ae. albopictus*, U4.4 cell line (SRR2182528) (Goic *et al*., 2016), whole mosquitoes (SRR2182437, SRR2182496) (Goic *et al*., 2016), female mosquito carcasses (SRR11095782, SRR11095783), ovary (SRR11095792, SRR11095793), and testes (SRR11252300) (Ma *et al*., 2021). For *Cx. quinquefasciatus*, Hsu cell line (SRR11252314) (Ma *et al*., 2021), whole mosquitoes (SRR5457467) (Dietrich *et al*., 2017), salivary glands (SRR8936292, SRR8936293, SRR8936298) (Rückert *et al*., 2019), midgut (SRR8936296, SRR8936297, SRR8936299) (Rückert *et al*., 2019), ovary (SRR11252316) (Ma *et al*., 2021), and testes (SRR11252317) (Ma *et al*., 2021). For *An. gambiae*, Mos55 cell line (SRR11252306) (Ma *et al*., 2021), head and thorax (SRR11713504, SRR11713505) (Bryant *et al*., 2020), midgut (SRR11713500, SRR11713501, SRR11713502) (Bryant *et al*., 2020), fat body (SRR11713503, SRR11713506, SRR11713507) (Bryant *et al*., 2020), ovary (SRR11713497, SRR11713498, SRR11713499) (Bryant *et al*., 2020), and testes (SRR11252311) (Ma *et al*., 2021). Different dataset from the same cell line or tissue were merged and used as a single library for downstream analysis.

After trimming of adaptors with Cutadapt (Martin, 2011), multi-mapping reads from the small RNA-seq libraries were randomly distributed over all possible mapping positions (--best –strata -M1) with Bowtie (Langmead *et al*., 2009). Annotation of piRNA clusters was performed as described (Crava *et al*., 2021; Palatini *et al*., 2020), with several parameters optimized (Figure S1A): 1) the minimum amount of piRNA-sized reads (23-33 nt) mapped to per 5-kb non-overlapping window, when normalized per million mapped piRNAs (ppm); 2) the uniquely mapped piRNA per cluster. A minimum of 20 ppm per 5-kb window and a minimum of 3 uniquely mapped piRNAs per cluster were used to annotate piRNA clusters from each cell line or tissue, separately. This piRNA cluster annotation pipeline was solely guided by piRNA coverage of the respective genomic regions, and did not include assumptions on nucleotide biases or strand asymmetry. In parallel, proTRAC was used to confirm piRNA cluster annotation, which requires strand asymmetry and enrichment for ping-pong signature (1U/10A) (Figure S1B) (Rosenkranz and Zischler, 2012).

#### Classification of core piRNA clusters and germline-specific piRNA clusters

Core piRNA clusters were defined as piRNA clusters that are universally expressed across cell lines and tissues. Within each mosquito species, we annotated piRNA clusters individually from each cell line/ tissue, and piRNA clusters that overlapped between at least six datasets are considered as core piRNA clusters. Bedtools intersect was used to retrieve overlapping piRNA clusters, and the boundaries of core piRNA clusters were based on cell line piRNA clusters. Besides, core piRNA clusters were also supported by clusters predicted by proTRAC (Rosenkranz and Zischler, 2012), exception *Cx. Quinquefasciatus*, due to the highly fragmented piRNA clusters predicted by proTRAC.

Germline-specific piRNA clusters were defined as piRNA clusters that are only expressed in germline tissues, ovary, and testes, but not in any other somatic tissues. Bedtools intersect was used to get overlapped piRNA clusters between ovary and testes, and with the option -v to remove piRNA clusters that overlapped with any cell line or somatic tissues (Quinlan and Hall, 2010). The boundaries of germline-specific piRNA clusters were based on ovary piRNA clusters. Detailed information of piRNA clusters of *Ae. aegypti* was summarized in Table S1, *Ae. albopictus* in Table S5, *Cx. quinquefasciatus* in Table S6, and *An. gambiae* in Table S7.

#### Characterization of piRNA clusters

piRNA cluster expression level was determined by mapping piRNA-sized reads (23-33 nt) to each piRNA cluster, using bamCoverage from deepTools (Ramírez *et al*., 2014). Pair-wise comparison of piRNA cluster expression was performed with log10 transformed piRNA cluster expression level using the R package ggplot2 (Wickham, 2016). Genome browser tracks of small RNA-seq datasets were generated with pyGenomeTracks from deepTools (Ramírez *et al*., 2014).

For transposable elements analyses, the repeat annotation files were downloaded for *Ae. aegypti* L5 (Vectorbase, Aedes-aegypti-LVP_AGWG_REPEATFEATURES_AaegL5.gff3), *Ae. albopictus* (NCBI, GCF_006496715.1), *Cx. quinquefasciatus*(Vectorbase, Culex-quinquefasciatus-Johannesburg_REPEATFEATURES_CpipJ2.gff3), and for *An. gambiae* (Vectorbase, Anopheles-gambiae-PEST_REPEATFEATURES_AgamP4.gff3). Bedtools were used to exact TEs that overlapped with either core piRNA clusters or germline-specific piRNA clusters.

Based on genomic distribution, a piRNA cluster is defined as a genic piRNA cluster, if the piRNA cluster is 80% covered in length by an expressed gene in the same orientation. Besides, a piRNA cluster is defined as a readthrough piRNA cluster, if *d* is below the threshold. We set a cutoff of *d* based on the genome size of each mosquito species, 5 kb for *Ae. aegypti*, 10 kb for *Ae. albopictus*, 2.5 kb for *Cx. quinquefasciatus*, and 1.0 kb for *An. gambiae*.

#### ChIP-seq and analysis pipeline

Aag2 cells were grown in Leibovitz’s L-15 medium (Invitrogen) supplemented with 10% fetal bovine serum (Gibco), 50 U/ml Penicillin, 50 μg/mL Streptomycin (Invitrogen), 1x Non-essential Amino Acids (Invitrogen) and 2% Tryptose phosphate broth solution (Sigma) at 28°C in a humidified atmosphere without CO_2_. Cell lines were maintained by splitting twice weekly according to confluency.

Aag2 cells were cross-linked with 1% formaldehyde for 10 min at room temperature with gentle shaking. Cross-linking was quenched with the addition of 1/10 volume 1.25 M glycine. Cells were washed with 1× PBS, then harvested by scraping in Buffer B (20 mM HEPES, 0.25% Triton X-100, 10 mM EDTA, and 0.5 mM EGTA). Cells were then pelleted by spinning at 600 × g for 5 min at 4 °C. Cell pellet was resuspended in Buffer C (150 mM NaCl, 50 mM HEPES, 1 mM EDTA, and 0.5 mM EGTA) and rotated for 10 min at 4 °C. Cells were again pelleted by spinning at 600 × g for 5 min at 4 °C. The cell pellet was resuspended in 1× incubation buffer (0.15% SDS, 1% Triton X-100, 150 mM NaCl, 1 mM EDTA, 0.5 mM EGTA, and 20 mM HEPES) at 15 million cells/mL. Cells were sheared with a Bioruptor Pico sonicator (Diagenode) at 4 °C using 6 cycles of 30 s on, 30 s off. Sonicated chromatin was spun at 18,000 × g for 5 min at 4 °C. Chromatin in the upper supernatant was aliquoted, immediately frozen in liquid nitrogen, and stored at −80°C.

Sonicated chromatin from 2 million cells was used for each ChIP experiment. Chromatin was incubated in 1× incubation buffer (0.15% SDS, 1% Triton X-100, 150 mM NaCl, 1 mM EDTA, 0.5 mM EGTA, and 20 mM HEPES) supplemented with 1× protease inhibitors, 0.1% BSA, and a 50:50 mix of Protein A and G Dynabeads (Invitrogen), with agitation overnight at 4 °C. Generally, 1 µg antibody was used for each ChIP (H3K4me3, Diagenode, C15410003; H3K27Ac, Diagenode, C15410196; H3K36me3, Diagenode, C15410058; H3K9me3, Diagenode, C15410193; RNA pol II, Santa cruz, sc-56767; RNA pol II ser5p, Abcam, ab5131; RNA pol II ser2p, Chromotek, 3E10). As a control, no antibody was added to the ChIP input sample (sonicated chromatin). The beads were washed twice with wash buffer 1 (0.1% SDS, 0.1% sodium deoxycholate, 1% Triton X-100, 150 mM NaCl, 1 mM EDTA, 0.5 mM EGTA, and 20 mM HEPES), once with wash buffer 2 (wash buffer 1 with 500 mM NaCl), once with wash buffer 3 (250 mM LiCl, 0.5% sodium deoxycholate, 0.5% NP-50, 1 mM EDTA, 0.5 mM EGTA, and 20 mM HEPES), and twice with wash buffer 4 (1 mM EDTA, 0.5 mM EGTA, and 20 mM HEPES). After the washing steps, beads were rotated for 20 min at room temperature in elution buffer (1% SDS and 0.1 M NaHCO_3_). The supernatant was de-cross-linked with 200 mM NaCl and 100 μg/mL proteinase K for 4 h at 65 °C. De-cross-linked DNA was purified with MinElute PCR Purification columns (Qiagen). DNA concentrations were determined by Qubit fluorometric quantification (ThermoFisher Scientific). As a quality check step, qPCR analysis of positive control and negative control regions using 10× diluted ChIP DNA was performed with the GoTaq qPCR system (Promega) according to the manufacturers’ recommendations. Detailed primer information of positive and negative controls is list in Table S2.

Libraries were prepared with the Kapa Hyper Prep Kit for Illumina sequencing (Kapa Biosystems) according to the manufacturer’s protocol with the following modifications. 3 ng DNA was used as input, with NEXTflex adapters (Bioo Scientific) and ten cycles of PCR amplification. Post-amplification cleanup and size selection was performed with dual AMPure XP beads selection (0.6× and 0.8×). Size-selected samples were analyzed for purity with a High Sensitivity DNA Chip on a Bioanalyzer 2100 system (Agilent), with an average size between 300 and 500 bp. Samples were sequenced in a paired-ended manner using the NextSeq 500 (Illumina) according to standard Illumina protocols.

After an initial quality check with FastQC (Andrews, 2010), sequencing reads were aligned to the *Ae. aegypti* reference genome (*Ae. aegypti* LVP, VectorBase release 56) using BWA (Li and Durbin, 2009). Duplicated reads were removed for further analysis. Mapped reads from two biological duplicates were merged and used for peak calling, because of good correlation (Pearson’s r > 0.98). Peak calling was performed with the MACS2 against an input sample with standard settings and a q value of 0.05 (Zhang *et al*., 2008), with the option of broad peak calling for H3K36me3 and H3K9me3. The peak files from four histone marks (H3K4me3, H3K27Ac, H3K36me3, and H3K9me3, detailed in Table S2) were used as input for chromatin state analysis using ChromHMM (Ernst and Kellis, 2017). The five-emission state model was chosen, as it represented biologically interpretable combinatorial patterns that were not repetitive or ambiguous and showed general agreement with previously published models (Gorkin *et al*., 2020; Kundaje et al., 2015; Yue et al., 2014) (Figure S2A). The deepTools package was used to generate bigwig files that are normalized to RPKM (Ramírez *et al*., 2014), and example track profiles were plotted with fluff (Georgiou and van Heeringen, 2016). Band plots and heatmaps were generated with computeMatrix and plotProfile from deepTools (Ramírez *et al*., 2014).

#### ATAC-seq and analysis pipeline

ATAC libraries were prepared essentially according to a protocol for mammalian cells with minor changes (Qu et al., 2019). In brief, Aag2 cells were trypsinized on plates and resuspended into single cells. Cells were pelleted by spinning at 500 × g for 5 min at 4°C and then washed twice with ice-cold 1× PBS buffer. 50,000 cells were counted and suspended in ice-cold 1× PBS. The same volume of 2× lysis buffer (20 mM Tris-HCl, 20 mM NaCl, 30 mM MgCl_2_, 0.5% IGEPAL) was added to 50,000 cells and pipetted up and down 20 times to lyse the cells. Cells were pelleted at 500 × g for 30 min at 4°C and suspended in 25 µL of digestion buffer (10 mM Tris pH 7.6, 5 mM MgCl_2_, 10% DMF, 1U Tn5 transposase), followed by tagmentation in the 37°C incubator for 1 h at 650 rpm. After tagmentation, 9 µL of clean-up buffer (0.5 M NaCl, 150 mM EDTA, 1% SDS, 2 mg/mL Protease K) was added for incubation at 40°C for 30 min with 650 rpm shaking. Normal phase 2× AMPure XP beads (68 µL) purification was performed to remove small fragments (<100 bp). Purified DNA was used for a first PCR amplification (Nextera Index Kit) with KAPA HiFi HotStart ReadyMix (Roche) for seven cycles. Reverse phase 0.55× SPRI bead (27.5 µL) purification was performed to remove large fragments (>300 bp), followed by purification with QIAquick PCR Purification Kit (Qiagen). Purified DNA was used for a second PCR amplification (Nextera Index Kit) with KAPA HiFi HotStart ReadyMix (Roche) for another seven cycles. The PCR product was purified again with QIAquick PCR Purification Kit (Qiagen). DNA amounts were determined with Qubit fluorometric quantification (ThermoFisher Scientific) and analyzed for fragment size distribution with a DNA 1000 Chip on a Bioanalyzer 2100 system (Agilent).

Samples were sequenced in a paired-ended manner using the NextSeq 500 (Illumina) according to standard Illumina protocols. After an initial quality check with FastQC (Andrews, 2010), sequencing reads were aligned to *Ae. aegypti* genome assembly (*Ae. aegypti* LVP, VectorBase release 56) using BWA (Li and Durbin, 2009). Potential PCR duplicates were removed using the Picard MarkDuplicates option. The filtered BAM files were used as input in MACS2 for peak calling with standard settings and a q value of 0.05 (Zhang *et al*., 2008) (Table S2). The deepTools package was used to generate, shift (alignmentSieve function), and normalize bigwig track files to RPKM (Ramírez *et al*., 2014), and example track profiles were plotted with fluff (Georgiou and van Heeringen, 2016). Band plots were generated with computeMatrix and plotProfile from deepTools (Ramírez *et al*., 2014).

#### Small RNA-seq and analysis pipeline

Small RNA deep sequencing libraries were generated using the NEBNext Small RNA Library Prep Set for Illumina (E7560, New England Biolabs), using 1 µg RNA as input. Library preparation was performed following the manufacturer’s instructions except for 3’ adapter ligation, which was performed for 18 h at 16°C to enhance the ligation efficiency of 2’-O-methylated small RNAs. Libraries were sequenced on an Illumina Hiseq 4000 machine by the GenomEast Platform (Strasbourg, France).

After an initial quality check with FastQC (Andrews, 2010), raw sequence reads were first trimmed with Cutadapt (Martin, 2011) and then mapped with Bowtie (Langmead *et al*., 2009). Reads in the size range from 21 to 24 bp were mapped to the pre-miRNA sequences of *Ae. aegypti* from miRBase database and annotated as miRNA (Kozomara et al., 2019). Reads in the size range from 25 to 30 bp were regarded as piRNAs. For PIWI-protein IP followed by small RNA-seq (PRJNA594491) (Halbach *et al*., 2020), piRNA read counts within a certain piRNA cluster were normalized to the corresponding IP input sample for enrichment. For analyzing the changes in piRNA expression after knockdown of genes of interest (Joosten *et al*., 2019; Joosten *et al*., 2021a; Joosten *et al*., 2021b; Miesen *et al*., 2015), piRNA reads counts were compared to the luciferase control knockdown after normalization to the total mapped miRNAs. Published small RNA-seq dataset used in this analysis include: knockdown of PIWI proteins (PRJNA266885) (Miesen *et al*., 2015), Veneno (SRR6395463-SRR6395468) (Joosten *et al*., 2019), Pasilla and Atari (PRJNA664195) (Joosten *et al*., 2021b), and Zuc (SRR12231665, SRR12231683, SRR12231685, SRR12231686, SRR12231693, SRR12231694) (Joosten *et al*., 2021a).

To analyze piRNAs mapped to nrEVEs, the latest nrEVEs annotations of *Ae. aegypti* (Crava *et al*., 2021) and *Ae. albopictus* (Palatini *et al*., 2020) were used. The length distribution was plotted in R based on the mapped reads ranging from 23 to 33 nt. For the analysis of nucleotide biases, piRNA reads were first separated based on the strand with SAMtools (Li *et al*., 2009b) and then trimmed to 20 nt for sequence logo representation with WebLogo (Crooks *et al*., 2004). Example tracks with nrEVE annotations were generated using pyGenomeTracks from the deepTools package(Ramírez *et al*., 2014).

#### RNA-seq analysis pipeline

Publicly available RNA-seq datasets were used to determine expressed genes in each mosquito species, including *Ae. aegypti* (SRR7938851, SRR7938854, SRR7938865) (Halbach *et al*., 2020), *Ae. albopictus* (SRR7907921, SRR7907923, SRR7907924), *Cx. quinquefasciatus* (SRR16931550, SRR16931551, SRR16931552), and for *An. gambiae* (SRR1525402, SRR1525403, SRR6127580, SRR6127581 (Pitts *et al*., 2014)). After initial quality control by FastQC (Andrews, 2010), raw sequence reads were aligned to the corresponding genome, *Ae. aegypti* LVP, *Ae. albopictus* Foshan FPA, *Cx. quinquefasciatus* Johannesburg, *An. gambiae* PEST (Vectorbase, release 56) using STAR (Dobin *et al*., 2013) with default settings. Quantification of gene expression (FPKM) was performed with Cuffdiff (Trapnell *et al*., 2013). Expressed genes were determined based on the distribution of FPKM per library (detailed in Table S3).

#### mRNA purification

Two rounds of mRNA purification were performed with NEB magnetic d(T)25 beads (S1419S, New England Biolabs), followed by DNaseI (Ambion)-treatment and reverse transcription, before cDNA PCR. PCR primers were listed in Table S4.

#### Stem-loop qPCR

Stem-loop qPCR (SL-qPCR) was performed essentially according to a protocol for miRNA quantification with minor changes (Chen et al., 2005). A total of 100 ng of RNA was reverse transcribed using Stem-loop reverse transcription primers (Table S4) and SuperScript II Reverse Transcriptase mix (Invitrogen). Stem-loop RT reaction conditions were as follows: 30 min at 16°C, 30 min at 42°C, and 5 min at 85°C. SYBR-green qPCR was performed as following: initial denaturation step for 5 min at 95°C, followed by 40 cycles of denaturation for 10s at 95°C, annealing for 20s at 60°C, and extension for 10s at 72°C. piRNA gene expression levels were normalized to the expression of the housekeeping miRNA Bantam and fold changes were calculated using the 2(-ΔΔCt) method (Livak and Schmittgen, 2001). All SL-qPCR primers were listed in Table S4.

#### RNAi screen in Aag2 cells

PCR products flanked by T7 promoter sequences at both ends were *in vitro* transcribed by T7 polymerase to produce dsRNA. Aag2 cells were seeded at a density of 1.5×10^5^ cells/well in a 24-well plate, 24 h before the dsRNA transfection in triplicates. For each target gene, a transfection mix containing 300 μl non-supplemented L-15 medium, 500 ng dsRNA, and 1.3 μl X-treme GENE HP DNA transfection reagent (Sigma) was prepared according to the manufacturer’s instructions. Per 24-well, 100 μl of the transfection mix was added in a dropwise manner. Three days post-transfection, Aag2 cells were lysed in 1 mL RNA-Solv reagent (Omega Bio-tek), followed by RNA extraction through phase separation and isopropanol precipitation. RNA integrity was evaluated on EtBr stained agarose gels, and RNA concentration was determined using the Nanodrop ND-1000.

For RT-qPCR analyses, DNaseI (Ambion)-treated RNA was reverse transcribed with random hexamers using the Taqman reverse transcriptase kit (Life Technologies) and PCR amplified using the GoTaq qPCR system (Promega) on a LightCycler 480 (Roche). In RNAi experiments, expression levels of target genes were normalized to the expression of the housekeeping gene lysosomal aspartic protease (*LAP*, *AAEL006169*), and fold changes were calculated using the 2-ΔΔCT method (Livak and Schmittgen, 2001). All primers were listed in Table S4.

## ACKNOWLEDGMENTS

We thank Chet H. Loh, Rebecca Halbach, Pascal Miesen, Ezgi Taşköprü, and members of the laboratory for fruitful discussion. This work is supported by a VICI grant from the Dutch Research Council (grant number 016.VICI.170.090).

## AUTHOR CONTRIBUTIONS

J.Q. and R.P.v.R designed the experiments and analyzed the data. J.Q. performed computational analyses and most of the experiments, with the help from V.B. and R.v.I. for the RNAi screen and F.M.K. for PCR validation of readthrough transcription. J.Q. and R.P.v.R. wrote the paper. All authors read and contributed to the manuscript.

## DECLARATION OF INTERESTS

The authors declare no competing interests.

## Extended data figures and tables

**Figure S1.**
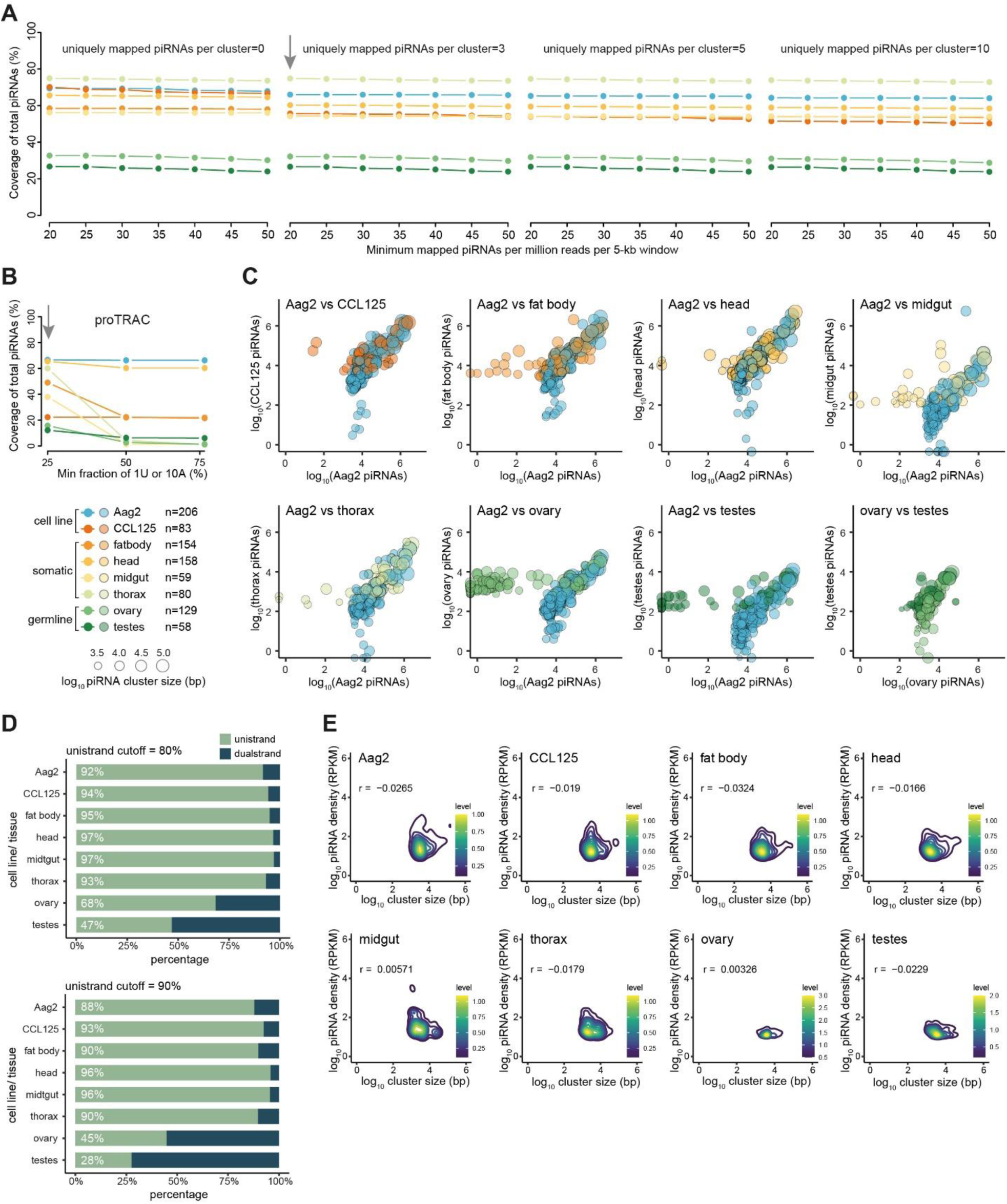
Annotation of piRNA clusters in *Ae. aegypti* mosquitoes. (A) Optimization of parameters for piRNA cluster prediction with an established pipeline that did not include assumptions on nucleotide biases or strand asymmetry (Crava *et al*., 2021; Palatini *et al*., 2020). Dot colors as in Figure 1A. (B) Optimization of parameters for piRNA cluster prediction with proTRAC (Rosenkranz and Zischler, 2012). Different thresholds of the ping-pong signature (1U or 10A) were tested. 25% was chosen for cluster prediction. (C) Comparison of piRNA expression of proTRAC-predicted piRNA clusters between cell lines, somatic and germline tissues. The same small RNA-seq datasets as in Figure 1A were used in this analysis. (D) Quantification of uni-strand and dual-strand clusters in the indicated cell lines and tissues, with an arbitrary threshold of 80% (top panel) or 90% (lower panel) piRNAs produced from one genomic strand defined as uni-strand piRNA clusters. (E) Correlation between piRNA cluster size and piRNA density, with Pearson’s r shown in the upper left corner and a level bar reflecting the number of clusters shown in the lower right corner.

**Figure S2.**
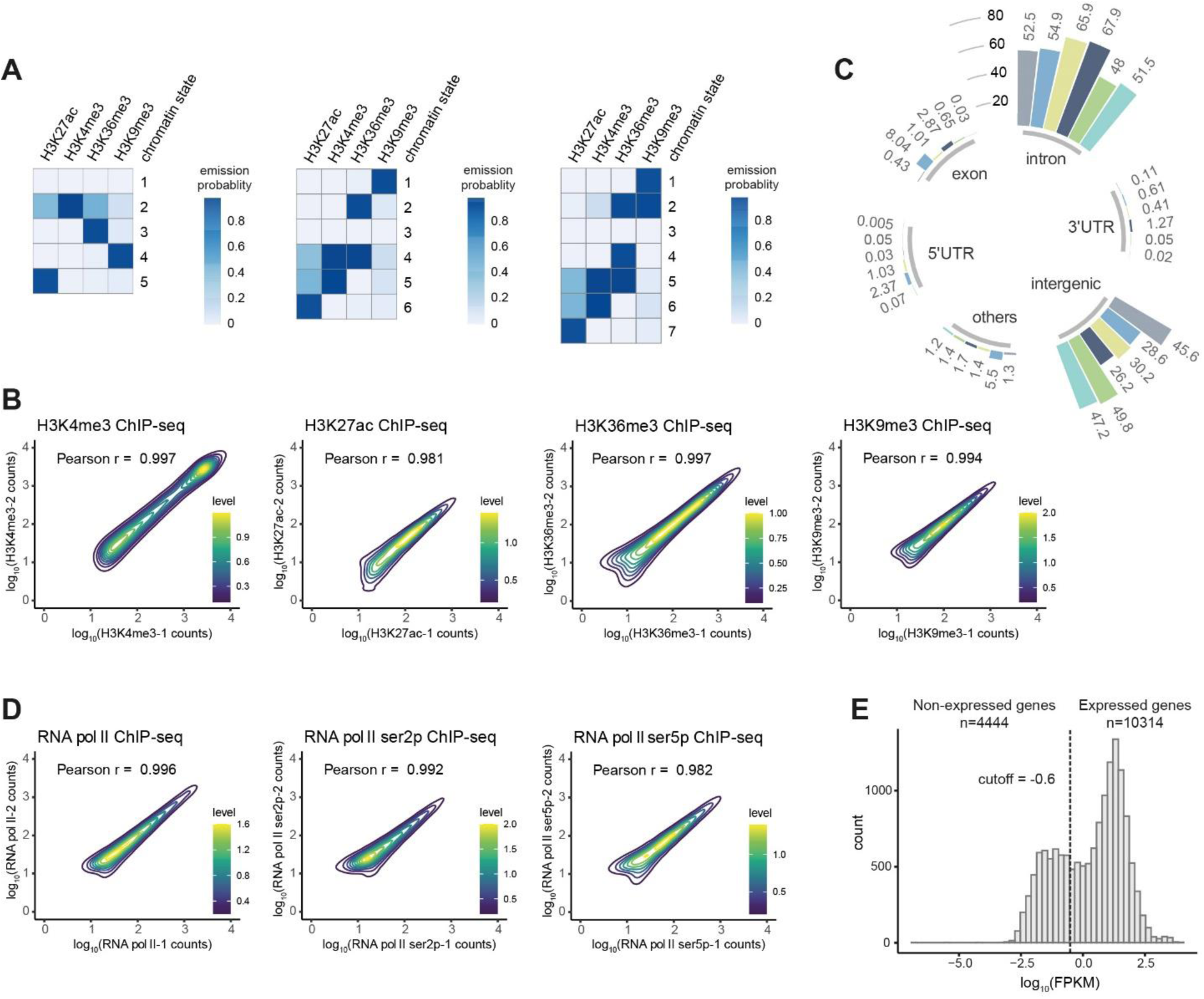
Quality check of the chromatin profiling dataset in Aag2 cells. (A) Chromatin state annotation using ChromHMM with different numbers of states based on ChIP-seq data in Aag2 cells. (B) Density profiles of ChIP-seq data of the indicated histone marks. Log scaled read counts within ChIP-seq peaks between two biological replicates were plotted. A level bar reflecting the number of ChIP-seq peaks is shown in the lower right corner. For detailed ChIP-seq peaks, see Table S2. (C) Genomic distribution of five chromatin states. Chromatin state colors are as in Figure 2A and the *Ae. aegypti* genome background distribution was shown as the 1^st^ gray bar. For instance, 54.9% of the promoter state was located in intron regions. The category ‘others’ consists of non-coding RNAs and pseudogenes. (D) Density profiles of RNA pol II, RNA pol II ser5p, and RNA pol II ser2p ChIP-seq in duplicates. For detailed information about peaks enriched for the three RNA pol II, see Table S2. (E) Published RNA-seq datasets were used to define expressed genes (Halbach *et al*., 2020). A cutoff of log_10_FPKM=-0.6 was chosen to distinguish expressed genes from non-expressed genes (Table S3).

**Figure S3.**
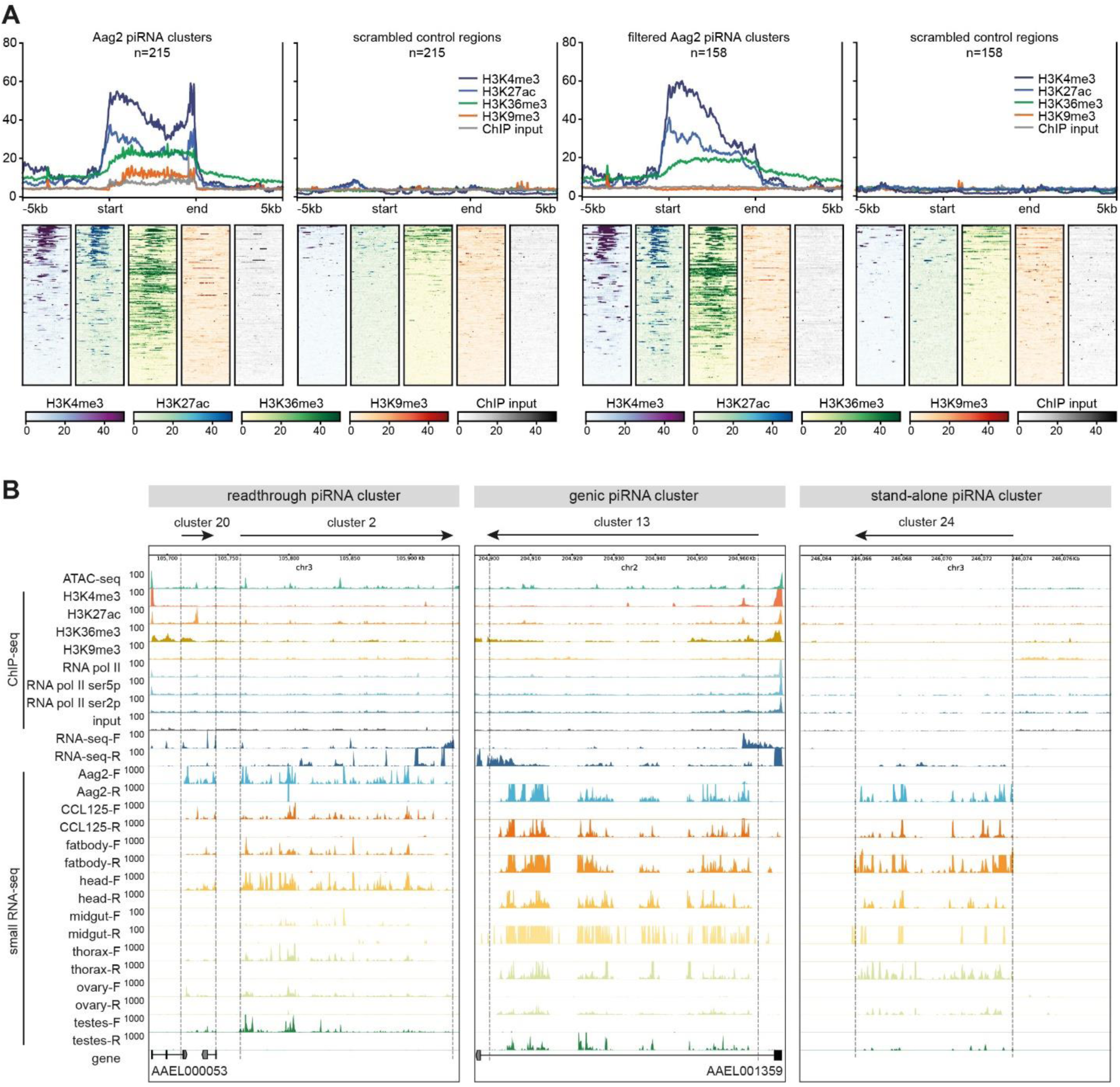
Chromatin signatures of Aag2 piRNA clusters. (A) Band plots (upper panel) and heat maps (lower panel) showing average signals of ChIP-seq data across 215 piRNA clusters in Aag2 cells (left) and 158 piRNA clusters (right) after removal of 57 piRNA clusters that overlapped with strong ChIP input signals (regions from MACS2 peak calling, described in M&M) to exclude background noise from genomic DNA. ChIP-seq signals were normalized to RPKM (y-axis). In the heatmap, all piRNA clusters were scaled to 5 kb, and the upstream/downstream 5 kb regions were shown. (B) Representative genome browser tracks showing examples of a readthrough piRNA cluster, a genic piRNA cluster, and a stand-alone piRNA cluster. ATAC-seq, ChIP-seq, and RNA-seq signals, as well as piRNA-sized reads from small RNA-seq, were normalized to RPKM. The boundaries of each cluster are indicated with dashed lines and the direction of piRNA clusters is indicated with arrows.

**Figure S4.**
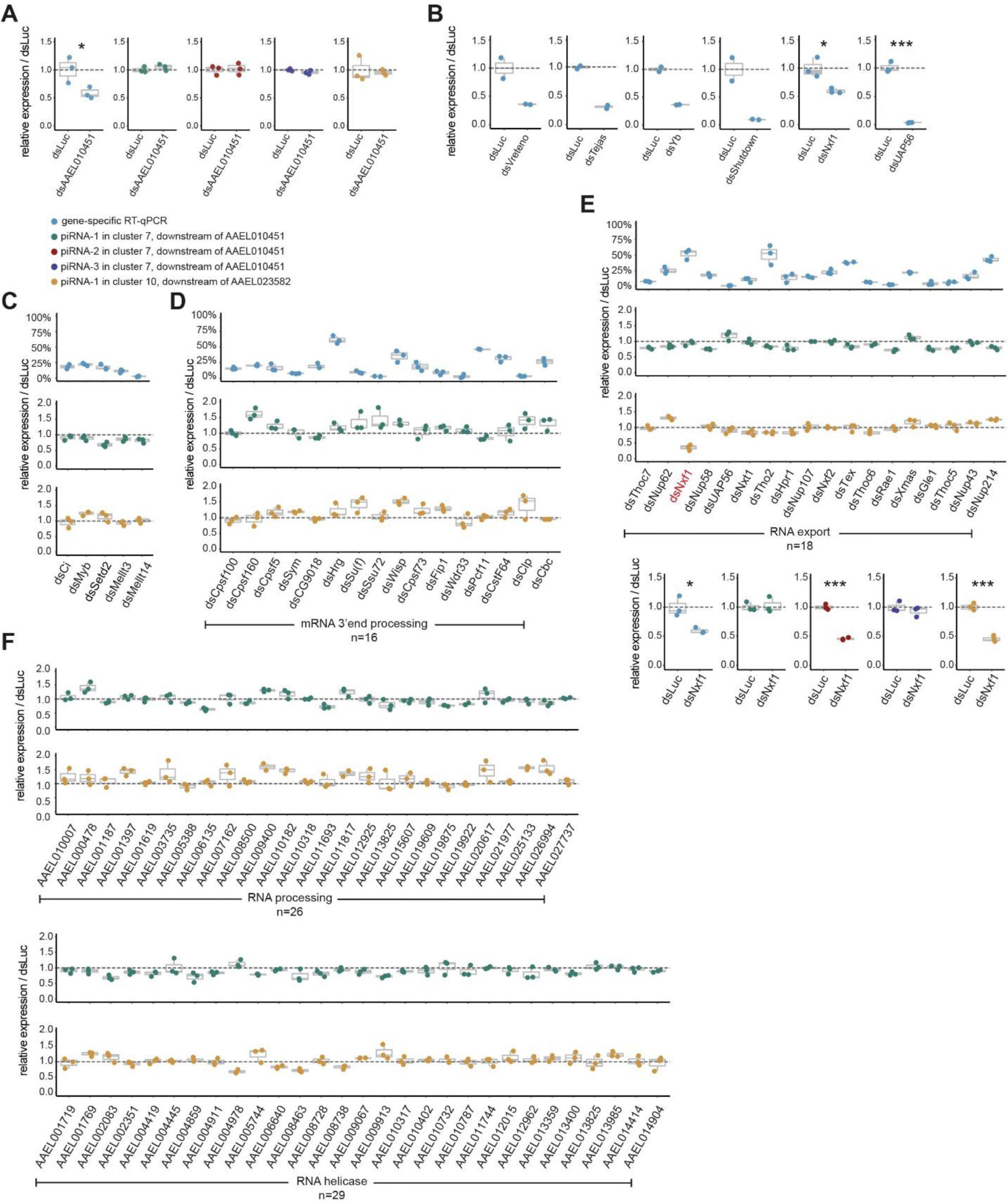
A large-scale RNAi screen to identify key factors involved in cluster-derived piRNA biogenesis. (A) Knockdown of *AAEL010451* using another dsRNA. RT-qPCR was performed to confirm gene knockdown, while SL-RT-qPCR was performed to quantify piRNA expression. All experiments were performed in biological triplicates (shown as dots) unless otherwise specified. The interquartile range and the median are shown in box plots. Statistical differences were examined by unpaired t-test, when compared to the non-targeting control knockdown of luciferase (dsLuc). Symbols denote statistical significance (* *p* < 0.05). For detailed primer information, see Table S4. (B) RT-qPCR was performed to confirm knockdown of target genes while SL-RT-qPCR was performed to quantify piRNA abundance. All experiments were performed in biological duplicates or triplicates (shown as dots). The interquartile range and the median were shown in box plots. Statistical differences were examined by unpaired t-test, when compared to the non-targeting control knockdown of luciferase (dsLuc). Symbols denote statistical significance (* *p* < 0.05, *** *p* < 0.001). For detailed primer information, see Table S4. (C)-(F) RNAi screen was performed with orthologs of known piRNA biogenesis factors in other organisms and chromatin modifiers. In other model systems, transcription factors such as A-MYB and Ci are important for piRNA cluster transcription (Goriaux *et al*., 2014; Li et al., 2013). We also tested the H3K36me3 writer, Setd2 (Sun et al., 2005), due to the observed readthrough transcription that was marked by rather a strong continuation of H3K36me3 signals (Figures 3A and D); and subsequently, two m^6A^ writers, Mellt3 and Mellt14, as recent studies suggested H3K36me3 guided the deposition of m^6A^ at 3’UTRs (Huang et al., 2019a), which may serve as a signal triggering readthrough transcription (C), 16 mRNA 3’end processing genes (D), 18 RNA export genes and confirmation of *Nxf1* knockdown (dsNxf1) with another set of dsRNA (lower panel) (E), 26 general RNA processing genes (Halbach *et al*., 2020) and 29 RNA helicase genes (Machado et al., 2022) (F).

**Figure S5.**
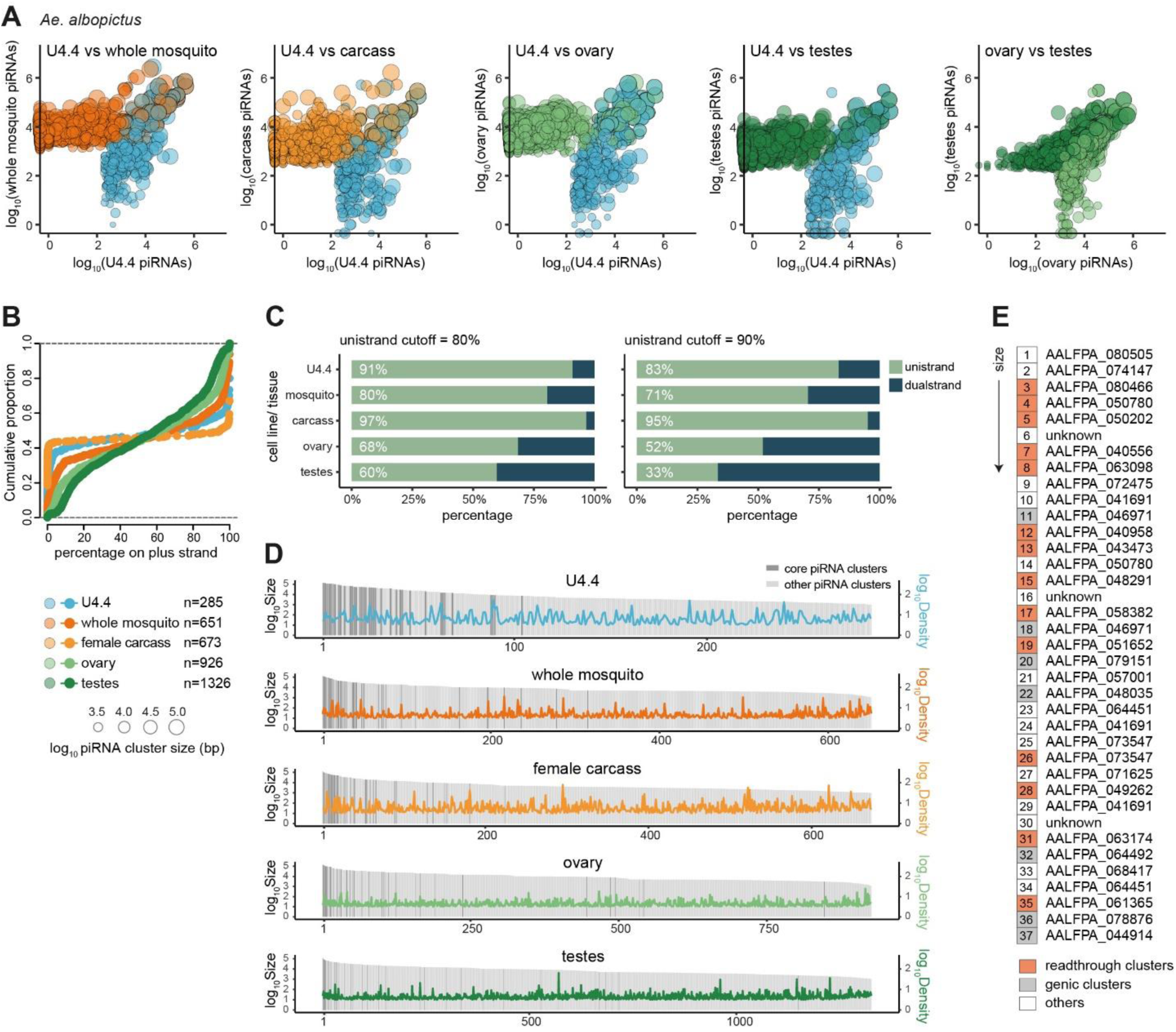
Systematic analysis of piRNA clusters in *Ae. albopictus* mosquitoes. (A) Comparison of piRNA expression within piRNA clusters between the U4.4 cell line, somatic tissue (carcass), and germline tissues (ovary and testes) of *Ae. albopictus*. piRNA clusters were independently annotated in each dataset and represented as dots, with color representing the origin and dot size representing cluster size. The numbers of clusters are listed. Log scaled piRNA read counts were used to represent piRNA expression levels within a piRNA cluster. For detailed piRNA cluster information, see Table S5. (B) Cumulative distribution of piRNA cluster coverage on the plus strand of the *Ae. albopictus* genome. (C) Quantification of uni-strand and dual-strand clusters in the indicated cell lines and tissues, with an arbitrary threshold of 80% or 90% piRNAs produced from one genomic strand defined as uni-strand piRNA clusters. (D) Distribution of 37 core piRNA clusters in cell lines and tissues. Within each dataset, piRNA clusters were ranked according to the cluster size. The 37 core piRNA clusters are shown as dark grey bars; other piRNA clusters are shown as light grey bars. piRNA density (RPKM) was plotted using the same colors as in (A). (E) Summary of the origin of core piRNA clusters, the distribution of nrEVEs, and the overlapping gene (for genic clusters) or the nearest expressed gene in the same orientation for readthrough piRNA clusters and other clusters. ‘Unknown’ refers to the situation when a piRNA cluster is located in a scaffold without any annotated gene nearby.

**Figure S6.**
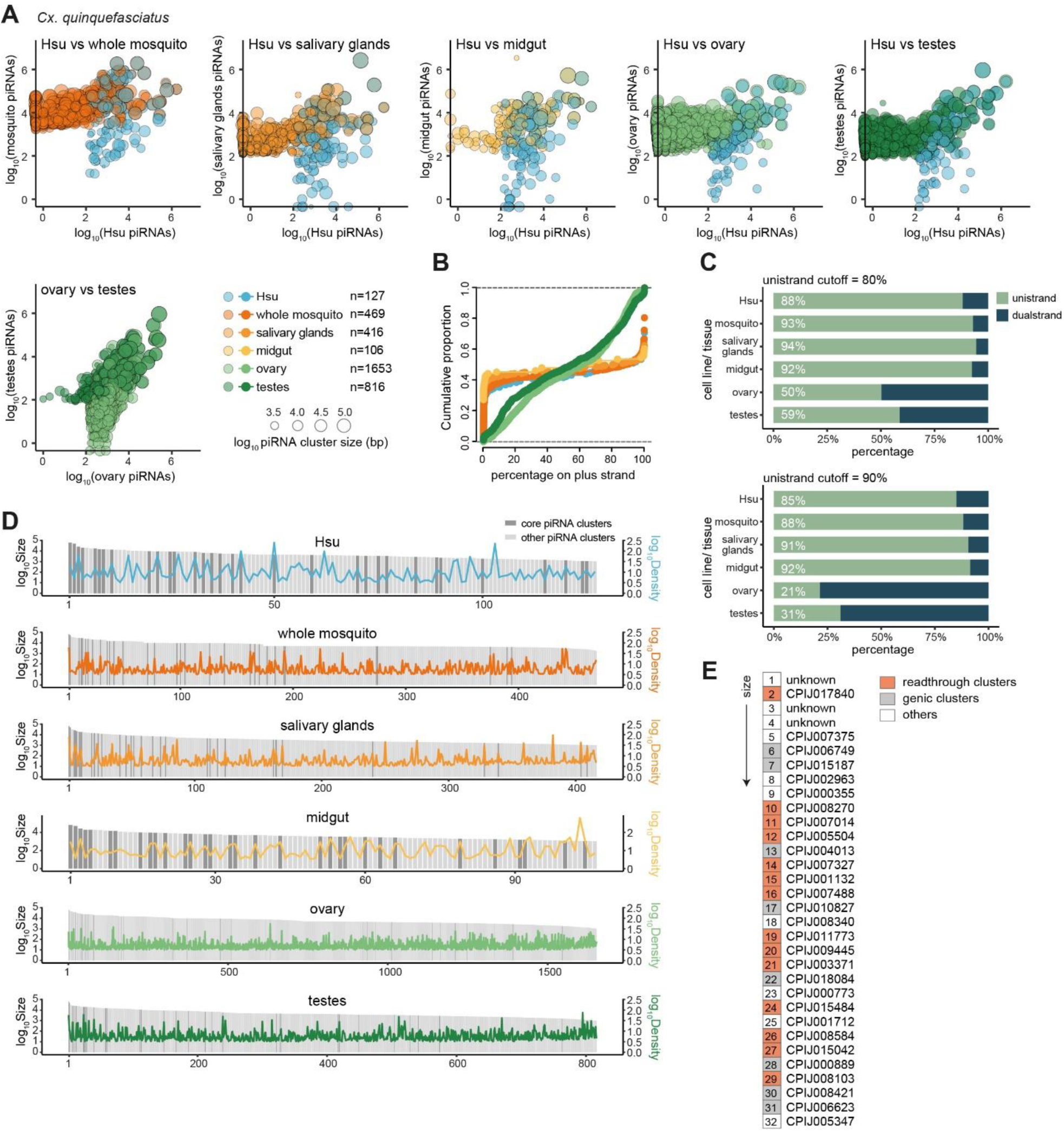
Systematic analysis of piRNA clusters in *Cx. quinquefasciatus* mosquitoes. (A) Comparison of piRNA expression within piRNA clusters between the Hsu cell line, somatic tissues (salivary glands and midgut), and germline tissues (ovary and testes) of *Cx. quinquefasciatus*. piRNA clusters were independently annotated in each dataset and represented as dots, with color representing the origin and dot size representing cluster size. The numbers of clusters are listed. Log scaled piRNA read counts were used to represent piRNA expression levels within a piRNA cluster. For detailed piRNA cluster information, see Table S6. (B) Cumulative distribution of piRNA cluster coverage on the plus strand of the *Cx. quinquefasciatus* genome. (C) Quantification of uni-strand and dual-strand clusters in the indicated cell lines and tissues, with an arbitrary threshold of 80% or 90% piRNAs produced from one genomic strand defined as uni-strand piRNA clusters. (D) Distribution of 32 core piRNA clusters in cell lines and tissues. Within each dataset, piRNA clusters were ranked according to the cluster size. The 32 core piRNA clusters are shown as dark grey bars; other piRNA clusters are shown as light grey bars. piRNA density (RPKM) was plotted using the same colors as in (A). (E) Summary of the origin of core piRNA clusters and the overlapping gene (for genic clusters) or the nearest expressed gene in the same orientation for readthrough piRNA clusters and other clusters. ‘Unknown’ refers to the situation when a piRNA cluster is located in a scaffold without any annotated gene nearby.

**Figure S7.**
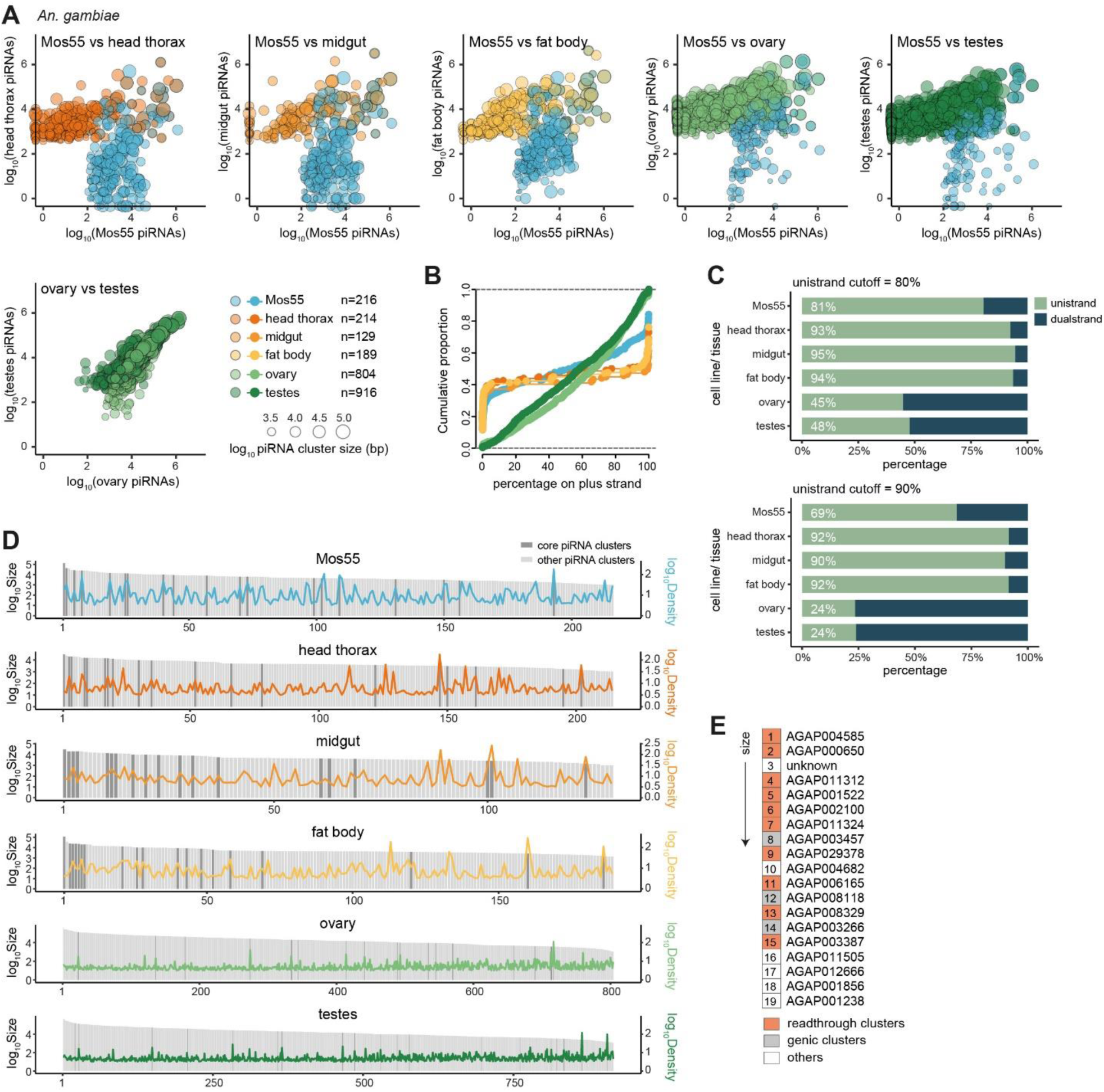
Systematic analysis of piRNA clusters in *An. gambiae* mosquitoes. (A) Comparison of piRNA expression within piRNA clusters between the Mos55 cell line, somatic tissues (head thorax, midgut and fat body), and germline tissues (ovary and testes) of *An. gambiae*. piRNA clusters were independently annotated in each dataset and represented as dots, with color representing the origin and dot size representing cluster size. The numbers of clusters are listed. Log scaled piRNA read counts were used to represent piRNA expression levels within a piRNA cluster. For detailed piRNA cluster information, see Table S7. (B) Cumulative distribution of piRNA cluster coverage on the plus strand of the *An. gambiae* genome. (C) Quantification of uni-strand and dual-strand clusters in the indicated cell lines and tissues, with an arbitrary threshold of 80% or 90% piRNAs produced from one genomic strand defined as uni-strand piRNA clusters. (D) Distribution of 19 core piRNA clusters in cell lines and tissues. Within each dataset, piRNA clusters were ranked according to the cluster size. The 19 core piRNA clusters are shown as dark grey bars; other piRNA clusters are shown as light grey bars. piRNA density (RPKM) is plotted using the same colors as in (A). (E) Summary of the origin of core piRNA clusters and the overlapping gene (for genic clusters) or the nearest expressed gene in the same orientation for readthrough piRNA clusters and other clusters. ‘Unknown’ refers to the situation when a piRNA cluster is located in a scaffold without any annotated gene nearby.

**Figure S8.**
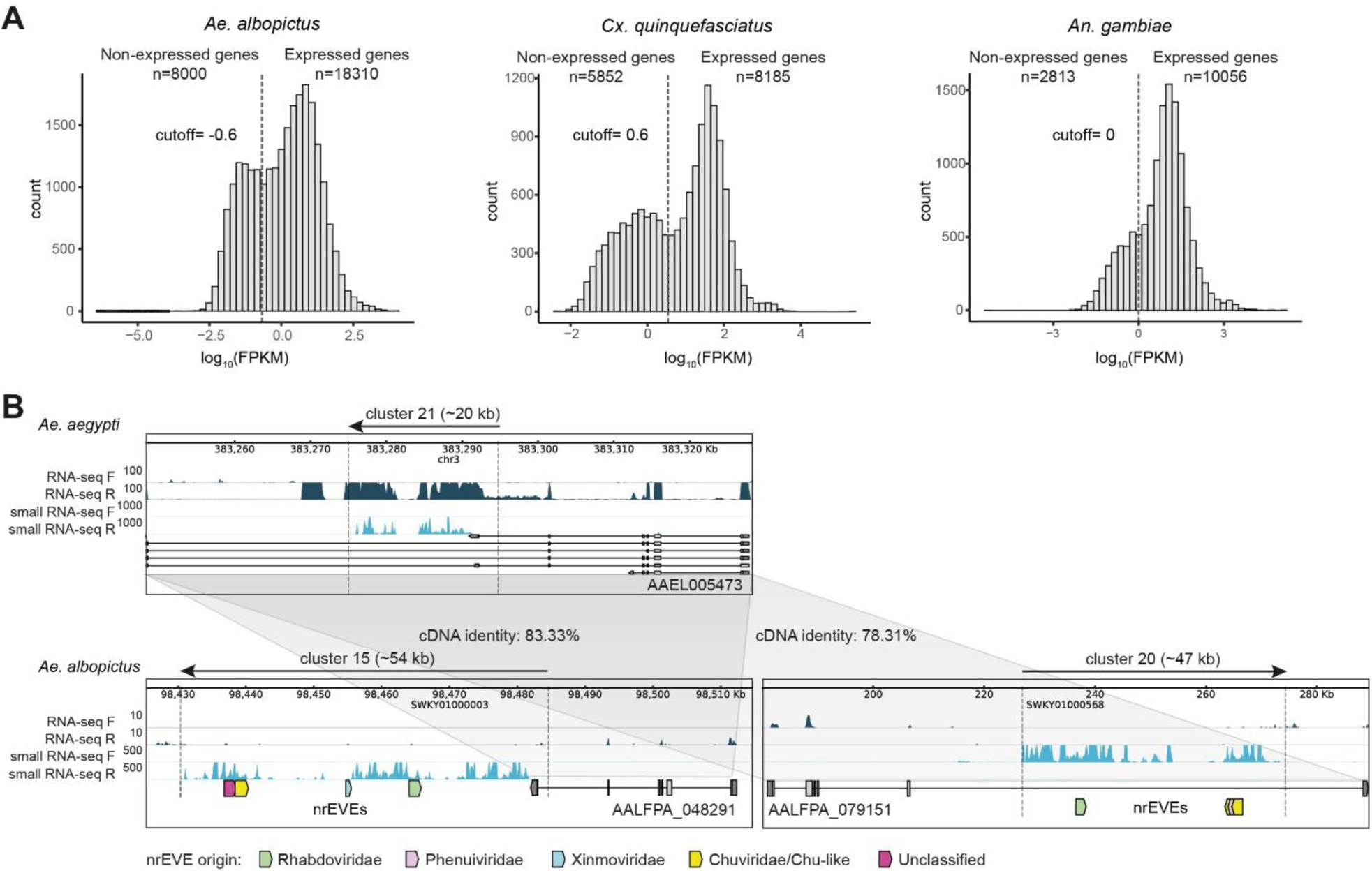
Comparison of upstream genes across different mosquito species. (A) Determination of the expressed genes in each species. A cutoff was chosen at the dip of the distribution plot to separate expressed genes from non-expressed genes, which are detailed in Table S3. (B) In *Ae. aegypti*, *AAEL005473* is associated with a genic piRNA cluster (cluster 21). Its two ortholog genes in *Ae. albopictus* are associated with a readthrough piRNA cluster (*AALFPA_048291*, cluster 15) and a genic piRNA cluster (*AALFPA_079151*, cluster 20). The boundaries of each cluster are indicated with dashed lines and the direction of the piRNA cluster is indicated with an arrow on top. nrEVEs were shown as arrows, with colors representing viral families.

## Supplemental Information

Figures S1-S8

Table S1. piRNA clusters in Ae. aegypti.xlsx

Table S2. chromatin profiles in Ae. aegypti Aag2 cells.xlsx

Table S3. Expressed genes in four mosquito species.xlsx

Table S4. A summary of primers.xlsx

Table S5. piRNA clusters in Ae. albopictus.xlsx

Table S6. piRNA clusters in Cx. quinquefasciatus.xlsx

Table S7. piRNA clusters in An. gambiae.xlsx

Table S8. Summary of nrEVEs in core piRNA clusters of Aedes mosquitoes.xlsx

## Notes

### Competing Interest Statement

The authors have declared no competing interest.

### Summary of Updates

Supplemental tables have been uploaded.

